# Distinct neural manifolds and critical roles of primate caudate nucleus in multimodal decision-making

**DOI:** 10.1101/2024.09.03.610907

**Authors:** Zhao Zeng, Ce Zhang, Yue Xu, Hua He, Yong Gu

## Abstract

Representational redundancy is ubiquitous in the brain with distributed networks encode task variables in a similar way, which may mask the unique roles of each region. Using a multisensory decision paradigm, here we showed that primate caudate nucleus (CN) dramatically differed from decision-related cortices in the reduced low-dimensional neural subspace, although possessing similar choice representations. Specifically, bimodal neural trajectories evolved towards nonvisual (vestibular) in CN, rather than towards visual as in frontal/parietal cortices. This challenges previous hypothesis that striatum mainly reflects cortical decision signals. Following recurrent neural networks simulations suggested that the cortico-subcortical distinction might be due to different intensities of single-modality inputs. We furtherly demonstrated that CN population responses represented the multisensory behavior strategy animal employed within the generalized drift-diffusion framework. Importantly, neural manipulations by unilateral drug injection and microstimulation confirmed CN’s causal contributions to multisensory decisions. Overall, our results indicate beyond relay-station in cortico-striatal circuitry, CN plays distinct and critical roles in perceptual decision-making.

## Introduction

Cognitive functions are mediated by neural networks in the brain, and it is frequent to see many nodes in the network share similar properties, leading to the so called “redundant” information processing hypothesis^1-3^. Perceptual decision making, for example, is one of the most important cognitive functions by which the brain transforms sensory inputs into perception, decision, and motor output^4^. During this process, ramping-like activity, reflecting evidence accumulation that leads to ultimate choice, has been observed widely distributed in both cortical and subcortical regions^3, 5, 6^ (Fig. 1a), including lateral intraparietal area^7^ (LIP), frontal eye field^8^ (FEF), superior colliculus^9^ (SC), caudate nucleus^10^ (CN). A key but unclear question is whether there is any inter-area difference in the choice representation. This is critical because the representational redundancy may mask the unique roles of individual regions^11^.

**Fig. 1.**
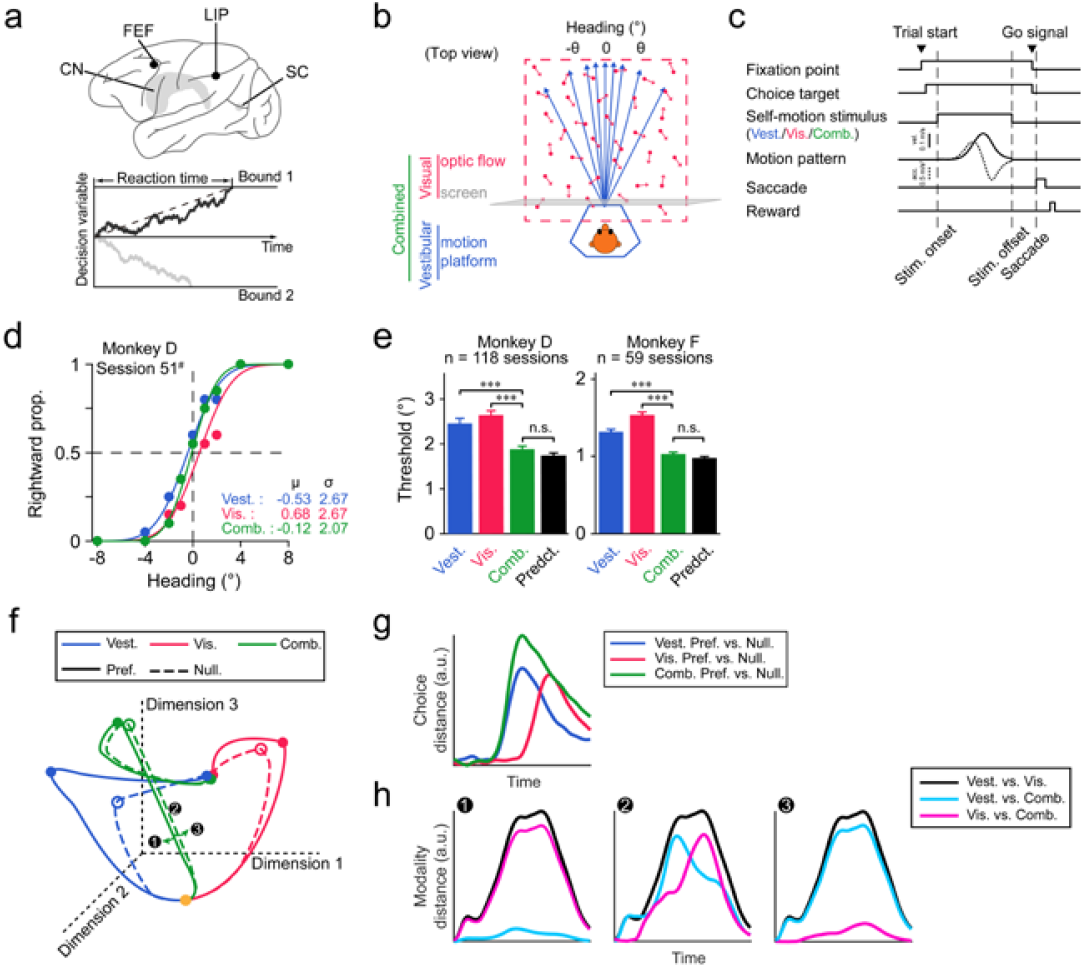
Comparing distributed perceptual decision-making signals using multisensory paradigm. **a**, Distributed brain regions (upper) representing evidence-accumulation signals (lower) in perceptual decision-making task. **b**, Schematic of a visuo-vestibular heading discrimination task using a virtual reality system. **c**, Timeline of a trial. Motion stimulus profile includes a varied Gaussian velocity (solid curve) and biphasic acceleration profile (dashed curve). **d**, Psychometric curves of an example session. The proportion of rightward choice is plotted as function of heading for each modality condition. Dots are real data, and curves are fitted cumulative Gaussian functions. **e**, Averaged psychometric thresholds of two monkeys for three modality conditions and for optimal prediction from Bayesian cue integration theory (black bar). Error bars indicate SEM; *** and n.s. respectively denote p < 1e-03 and no significance. **f**, Conceptual illustration of possible population activity projected onto a three-dimensional subspace during multisensory decision-making task. Color: modality (blue, vestibular-only; red, visual-only; green, cue-combined); Solid and dashed lines: animal choice in the preferred target (IN) and anti-preferred target (OUT). Orange dot: start point of a trial. **g**, Hypothetical choice-related signals quantified by Euclidean distance between IN and OUT trajectories in the subspace of **f**. **h**, Hypothetical modality-related signals quantified by Euclidean distance between neural trajectories across modalities in the subspace of **f**. Three possible models were proposed for the bimodal neural trajectory: biased towards vestibular-only condition (left panel, model #1), balanced in the middle of the two single cues (middle panel, model #2), or biased towards visual-only condition (right panel, model #3). Color represents comparison between different modalities: (black, vestibular-only versus visual-only; cyan, bimodal versus vestibular-only; magenta, bimodal versus visual-only).

Among these areas, the dorsal medial striatum (DMS), named as caudate nucleus (CN) in primates, is well known for mediating a variety of brain functions, including motor execution^12, 13^, value evaluation^14, 15^, time perception^16, 17^ and more. What is the role of CN in the distributed sensory-motor association network? A general hypothesis is that choice signals in CN largely stem from cortical regions such as LIP and FEF^18, 19^. Recent works even suggest that caudate mainly reflects cortical activity^20, 21^, serving as a relay station for the cortico-striatal circuitry. This hypothesis makes sense in term of that CN receives heavy projections from the neocortex^22^. yet it is inconsistent with the finding that the causal role of DMS but not those cortical decision areas has been confirmed^23-25^.

In the current study, we addressed these issues by using a multisensory decision-making paradigm, in which an additional dimension of sensory modality was provided. Specifically, macaques were trained to discriminate heading directions based on reliability-matching visual (optic flow) and vestibular (inertial motion) cue (Fig. 1b,c). After training, the animals showed roughly equal performance in single cue conditions, as reflected by similar psychophysical threshold (Fig. 1d,e). The threshold was significantly reduced in bimodal condition when congruent visuo-vestibular cues were provided, and the performance improvement was close to Bayesian prediction, a pattern typically seen in previous studies^26-28^. During the task, neural activity of single-units from CN, FEF, and LIP were recorded. In population, we expect manifold subspace reduced from high-dimension neural state space (Fig. 1f), would exhibit two properties about choice and modality. First, choice-related signals should evolve, reflecting momentary sensory evidence accumulation e.g. visual velocity and vestibular acceleration^27, 28^, a process which ultimately leads to different decisions (Fig. 1g). Second, modality-related signals should evolve and diverge for each single cue conditions (e.g. visual and vestibular). As to bi-modal stimulus condition, neural trajectory is speculated to be in the middle of the two single cues who share similar reliabilities (Fig. 1f), which can be quantified by a Euclidean distance index among modalities (Fig. 1h, middle panel, model #2). Alternatively, the bimodal neural trajectory may be biased towards either cue (Fig. 1h, first and third panel, model #1 and #3, respectively). We then examine whether the distributed networks would present similar or distinctive patterns along these two main dimensions (choice and modality).

## Results

### CN encodes multiple variables in multimodal decision-making task

As briefly aforementioned, two macaque monkeys (monkey D and F) were trained to perform a visuo-vestibular heading discrimination task under a fixed-duration (FD) two-alternative forced choice paradigm (Fig. 1b). In each trial, a linear forward heading stimulus with a small deviation from straight ahead was offered, lasting 1500 ms. After a random delay (300-500 ms), the monkeys were required to report their perceived direction, either left or right, by making a saccade to the corresponding target to earn reward (Fig. 1c). There were three sensory modality contexts: 1) vestibular-only condition, in which heading stimulus was provided solely by physical movement of a motion platform; 2) visual-only condition, in which heading was only simulated by optic flow on a display; 3) cue-combined condition, in which congruent vestibular and visual stimuli were offered at the same time. Note that cue-reliability was carefully manipulated so that the behavioral performance would be similar between the two single cue conditions^26^. All stimuli conditions, including three sensory modalities and nine heading angles, were interleaved in one experimental session. After training, both monkeys performed well, illustrated by nice psychometric curves (Fig. 1d). To quantify the performance, we fitted the psychometric curves via cumulative Gaussian function and took the parameters of mean (μ) and standard deviation (σ) to represent point of subjective equality (PSE) and psychophysical threshold, respectively. It turned out in both animals, psychophysical threshold was comparable in the visual-only and vestibular-only condition, and became significantly smaller in the cue-combined condition (Fig. 1e). Importantly, performance under bimodal condition was close to that predicted from Bayesian optimal integration theory (Fig. 1e, green vs. black bar), which has been frequently observed in many previous studies^26-29^.

While the animals performed the task, 200 well-isolated neurons were recorded from CN. Among them, 169 (monkey D, 117; monkey F, 52) putative medium spiny neurons (MSN) were identified according to their electrophysiological properties (Extended Data Fig. 1a-c)^30^. Peristimulus time histograms (PSTHs) of a few example neurons were shown to demonstrate that firing patterns of single CN neurons contained large heterogeneity (Extended Data Fig. 1d). To quantify the strength of task variables, we computed choice divergence (CDiv, Extended Data Fig. 1e) and modality divergence (MDiv) for each neuron, using receiver operator characteristic (ROC) analysis. For CDiv, many CN neurons significantly possessed choice signals in at least one modality condition (monkey D, 60/117 = 51.3%; monkey F, 33/52 = 63.5%, Extended Data Fig. 3e). The mean CDiv was significantly larger than 0 (vestibular-only: 0.123, p = 9.75e-06; visual-only: 0.099, p = 2.83e-04; combined: 0.139, p = 5.28e-06; two-tailed t-test; Fig. 2a), indicating that CN overall prefers contralateral choice. For MDiv, many CN neurons also carried significant sensory modality signals (monkey D: 88/117 = 75.2%; monkey F: 36/52 = 69.2%), with roughly equal visual and vestibular preference (mean MDiv: -0.012, p = 0.717; two-tailed t-test; Fig. 2a). These cells with choice or modality preference were distributed along the longitudinal axes in a mixed manner (Extended Data Fig. 1f,g).

**Fig. 2.**
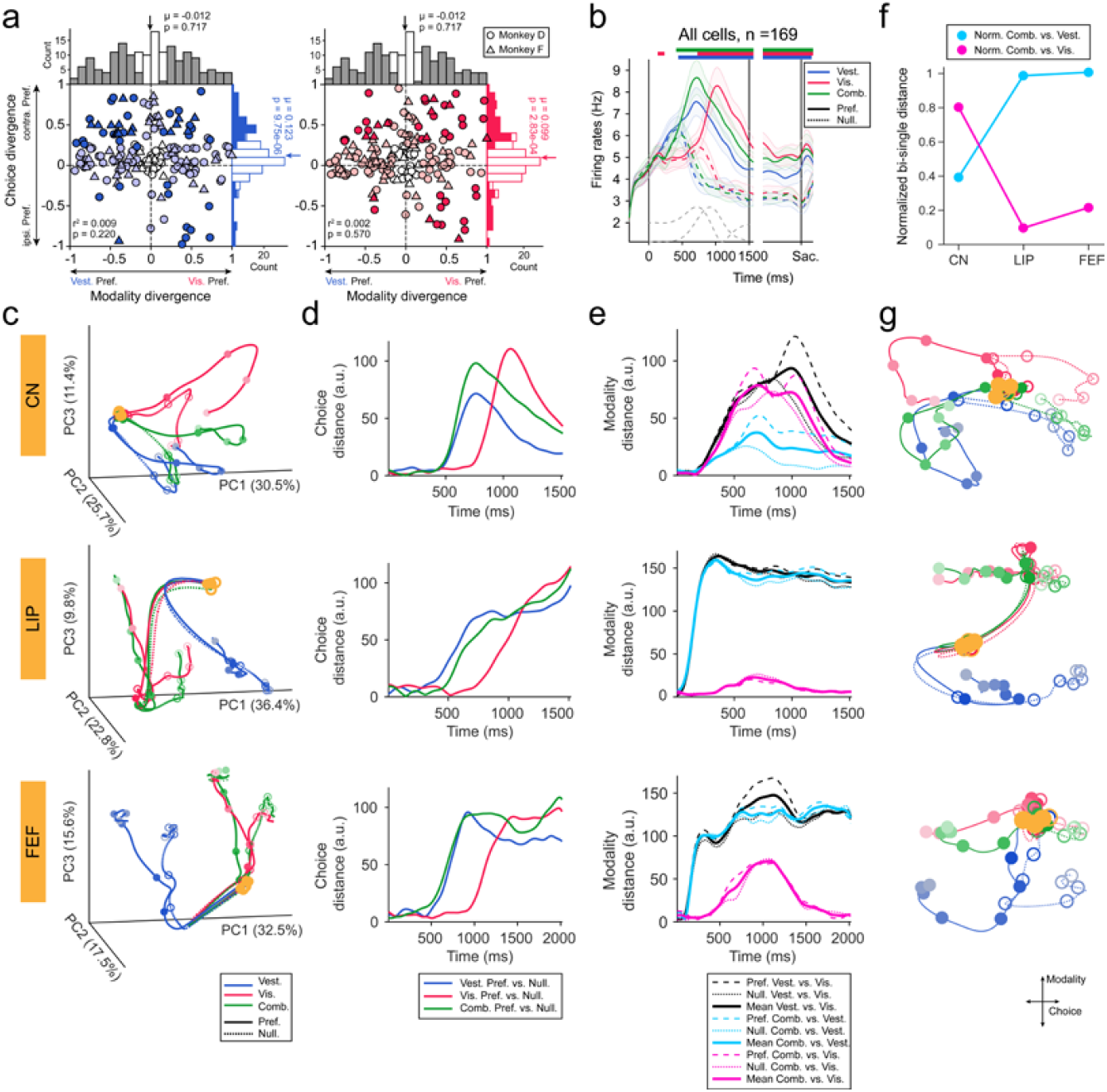
CN manifold and comparison with frontal and parietal association cortices. **a**, Choice and modality signals in each CN neuron in vestibular-only (left) and visual-only (right) conditions. Dark dot: both choice and modality signals are significant; Light dot: either variable is significant; Empty dot: neither variable is significant (p<0.01; permutation test). Shape of dot: different monkeys. **b**, Averaged population PSTHs of CN neurons. Firing rates are aligned to stimulus onset (1^st^ vertical line) and offset (2^nd^ vertical line), or saccade onset (3^rd^ vertical line). Shaded error bars are SEM. Horizontal colored bars indicate significant choice divergence (p < 0.01, two-tailed t-test). **c**, High-dimensional neural state space in CN (upper), LIP (middle), and FEF (lower) are projected onto 3-dimensional subspace constructed by the first three principal components. All labels are same as Fig. 1f. **d**, Choice distance evolved within trials in CN (upper), LIP (middle), and FEF (lower). All labels are same as Fig. 1g. **e**, Modality distance evolved within trials in CN (upper), LIP (middle), and FEF (lower). Thin dashed and dot lines respectively represent modality distance of IN and OUT trials, while thick solid lines are mean value of both IN and OUT trials. **f**, Normalized modality Euclidean distance computed by averaged modality distance across time for the bimodal versus unimodal conditions (cyan or magenta curves in E), normalized by modality distance of two uni-modal conditions (black curve in E). **g**, High-dimensional neural activities in CN (left), LIP (middle), and FEF (right) are projected onto subspace spanned by the two orthogonal axes of choice and modality using TDR analysis.

Previous studies have shown that sensory-motor association areas such as the posterior parietal cortex encodes multiple task variables using a manner of category-free coding in multisensory decision tasks^27, 31^. This was the similar case in CN. Firstly, numerous CN cells simultaneously had choice and modality preferences (vestibular-only: 47/169 = 27.8%; visual-only: 43/169 = 25.4%; combined: 54/169 = 31.9%). Secondly, there were neither well-clustered distribution nor significant correlation between choice and modality signals (Fig. 2a). However, either task variable could be successfully extracted from CN population activity by using linear demixing algorithms such as demixed principal component analysis (dPCA, Extended Data Fig. 1h)^32^. We found the interaction term of choice and modality also captured response variances to some extent (Extended Data Fig. 1h, the third column). This nonlinear mixed selectivity has been previously shown to be more computationally flexible for dealing with complex cognitive tasks^33^.

At the population level, we sorted trials in each modality condition according to animal choices and averaged firing rates of all neurons to construct population PSTH, finding significant choice modulation under all three contexts (Fig. 2b). Similar to LIP/FEF^27, 28^, choice-related signals in visual-only context were delayed compared to the other two stimulus conditions, implying that CN also integrates different dynamic signals, that is, vestibular acceleration but visual velocity (Extended Data Fig. 1i and Extended Data Fig. 2a). Majority of CN neurons are tuned to subjects’ internal choices but not external physical heading stimuli, which was supported by similar firing pattern in correct and error trials (Extended Data Fig. 3a,b) and markedly larger partial correlation between choice and CN neuronal responses (Extended Data Fig. 3c). More directly, we tested heading tuning of CN single neurons in the passive task where decision reports were not required, and found that very limited proportion of CN neurons were significantly selective to sensory headings (Extended Data Fig. 3d). Besides, choice signals for each neuron were highly correlated across modalities (Extended Data Fig. 3f).

In summary, these basic neural properties indicate that CN is within the sensory-motor association network, by showing similar patterns at the first glance as seen in other association cortices such as LIP and FEF.

### CN shows distinct neural manifold from frontal and parietal association cortex

Sensory-motor association areas usually contain dynamic and mixed signals that are not easily identified on single neurons, thus we constructed high-dimensional neural state space^34^ for population activity in each area including CN, FEF and LIP, and then performed dimension reduction to capture principal traits. Specifically, population activity in each region was denoised by using principal component analysis (PCA), and the first three PCs (captured more than 65% variances) were adopted to form a 3-D neural subspace (Fig. 2c). Two features could be readily extracted from this manifold subspace. First, neural trajectories sorted by choice evolved from a common starting point, and then gradually diverged in all three stimulus conditions as a function of time. Such a pattern is qualitatively similar in all areas. Indeed, Euclidean distance between each pair of trajectories, defining choice distance (Fig. 2d) illustrated a ramp-like trait with an obvious visual delay compared to vestibular in all three areas. Second, in contrast to the choice dimension, we found that neural trajectories sorted by modality dramatically differed across areas. Specifically, neural trajectory in the bimodal condition was fairly close to vestibular in CN, while was on contrary close to visual in LIP and FEF, as clearly seen in the manifold subspace (Fig. 2c), as well as the quantified Euclidean distance between each pair of modality comparison (Fig. 2e,f). All these results were further confirmed by a targeted dimensionality reduction (TDR) algorithm^35^, by which the high-dimensional neural population activities were directly projected onto a meaningful subspace spanned by choice and modality axes (Fig. 2g). These results also held across animals (Extended Data Fig. 4).

Thus, along modality dimension, CN manifold exhibits a distinct pattern from the frontal and posterior-parietal cortices. In fact, none of the association areas shows the expected symmetric pattern (model #2 in Fig. 1h), which is surprising because the two single cue reliabilities have been carefully matched in our study, as evident from two observations. First, in behavior, all animals showed analogous psychophysical threshold in each single cue condition (Fig. 1d,e). Second, in neurophysiology, one earlier stage polysensory area─the dorsal medial superior temporal sulcus (MSTd) in fact exhibited the expected model #2 pattern: bimodal neural trajectory was in the middle of the two single cues (Extended Data Fig. 5). Thus, CN may have received vestibular signals elsewhere in addition to FEF/LIP/MSTd, leading to an internally higher vestibular “gain” that attracts the neural trajectory towards vestibular during bimodal stimulus condition.

### Recurrent neural network reproduced inter-area neural state heterogeneity

To help understand possible sources of inter-area heterogeneity in neural state trajectory along modality dimension, we turned to a recurrent neural network (RNN) constituting fully connected, nonlinear neurons who performed a visuo-vestibular heading discrimination task, in a way similar to the animals. Specifically, RNN neurons received noisy heading evidence from vestibular acceleration and visual velocity inputs (Fig. 3a), and were connected to a single linear read-out decoder, who made decisions based on all hidden units’ activities via weighted summation. In each simulation, the RNN’s only goal was to categorize heading directions either leftward or rightward as accurately as possible, based on vestibular-only, visual-only, or bimodal sensory inputs. During training, the network initially showed relatively high thresholds, yet they decreased soon after tens of trials before reaching a plateau (Fig. 3b,e,h, middle panels). Interestingly, the network automatically showed improved performance under bimodal stimulus condition, close to the Bayesian prediction as seen in the animals’ behavior (Fig. 3b,e,h, right panels).

**Fig. 3.**
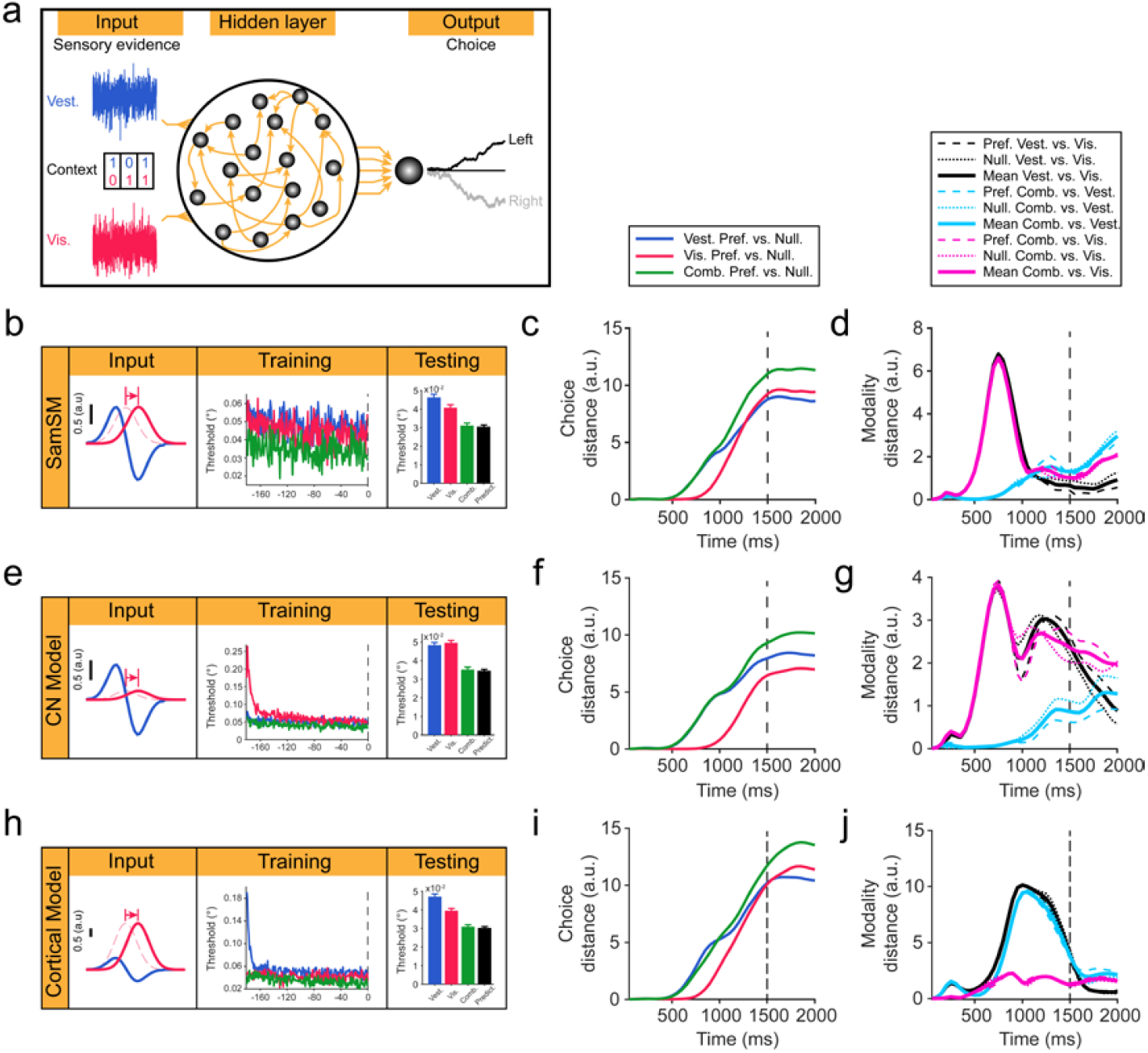
RNN simulations reproduced inter-area heterogeneity. **a**, Structure of a recurrently connected artificial network constituting 256 nonlinear units receiving noisy vestibular and visual heading inputs. **b-d**, Comparable vestibular and visual input magnitude and signal-to-noise ratio (**b**, left panel). RNN output behavioral performance (**b**, mid and right panels), choice (**c**) and modality (**d**) patterns. Vertical dashed line in (**c**) and (**d**) represents stimulus offset. **e-g**, Higher vestibular input magnitude with comparable signal-to-noise ratio (**e**, left panel). RNN output behavioral performance (**e**, mid and right panels), choice (**f**) and modality (**g**) patterns. **h-j**, Higher visual input magnitude with comparable signal-to-noise ratio (H, left panel). RNN output behavioral performance (**h**, mid and right panels), choice (**i**) and modality (**j**) patterns.

We then intuitively varied the magnitude ratio of the two unisensory inputs fed into the network (Fig. 3b,e,h, left panels). Importantly, noise level was always controlled so that signal-to-noise ratio of either input remained unchanged, to guarantee output of analogous visual and vestibular performance for each single cue (Fig. 3b,e,h, right panels). However, we hoped that the different magnitude factor, might represent some different impacts (e.g. over- or under-weighting) on particular cues during bimodal stimulus condition. This was indeed the case for our RNN simulation. In particular, RNN’s bimodal state trajectory evolved towards vestibular when vestibular has an equal (Fig. 3d) or relatively larger magnitude (Fig. 3g) compared to visual, generating a CN-like pattern. On contrary, only when visual has a much larger magnitude (Fig. 3j), would RNN produce a frontal/parietal-like visual bias during bimodal stimulus condition.

In contrast to modality dimension, RNN produced similar choice patterns that were independent on input magnitude-ratio (Fig. 3c,f,i). Therefore, RNN results could largely reproduce neurophysiological findings across cortico-subcortical areas. The heterogeneity in modality dimension across areas though, may suggest that CN receives different vestibular inputs other than simply inheriting everything from frontal and posterior-parietal association cortices.

### CN dynamics reflect perceptual performance in multisensory context

Not reflecting cortical decision representation, will CN play critical roles in the multimodal heading discrimination task per se? We first addressed how CN neuronal dynamics were functionally coupled with the animals’ perceptual performance on a single-trial basis using a number of analytical methods, and then examined whether such correlations were “causal” using perturbation methods as would be illustrated in the next sections.

The first analysis was to train a series of Lasso decoders over time to predict animal’s choice on a single-trial basis from CN’s population activity (see Method). We found monkeys’ upcoming choice could be reliably decoded about 500 ms after stimulus onset in the vestibular-only and bimodal conditions, and about 850 ms in visual-only condition (Fig. 4a, principal diagonal). Importantly, classifiers trained using activity during early period could be generalized to later stage both within, and across stimulus conditions (Fig. 4a, off the principal diagonal), implying that choice signals in CN share similar strategy across time and modalities (e.g. abstract decision).

**Fig. 4.**
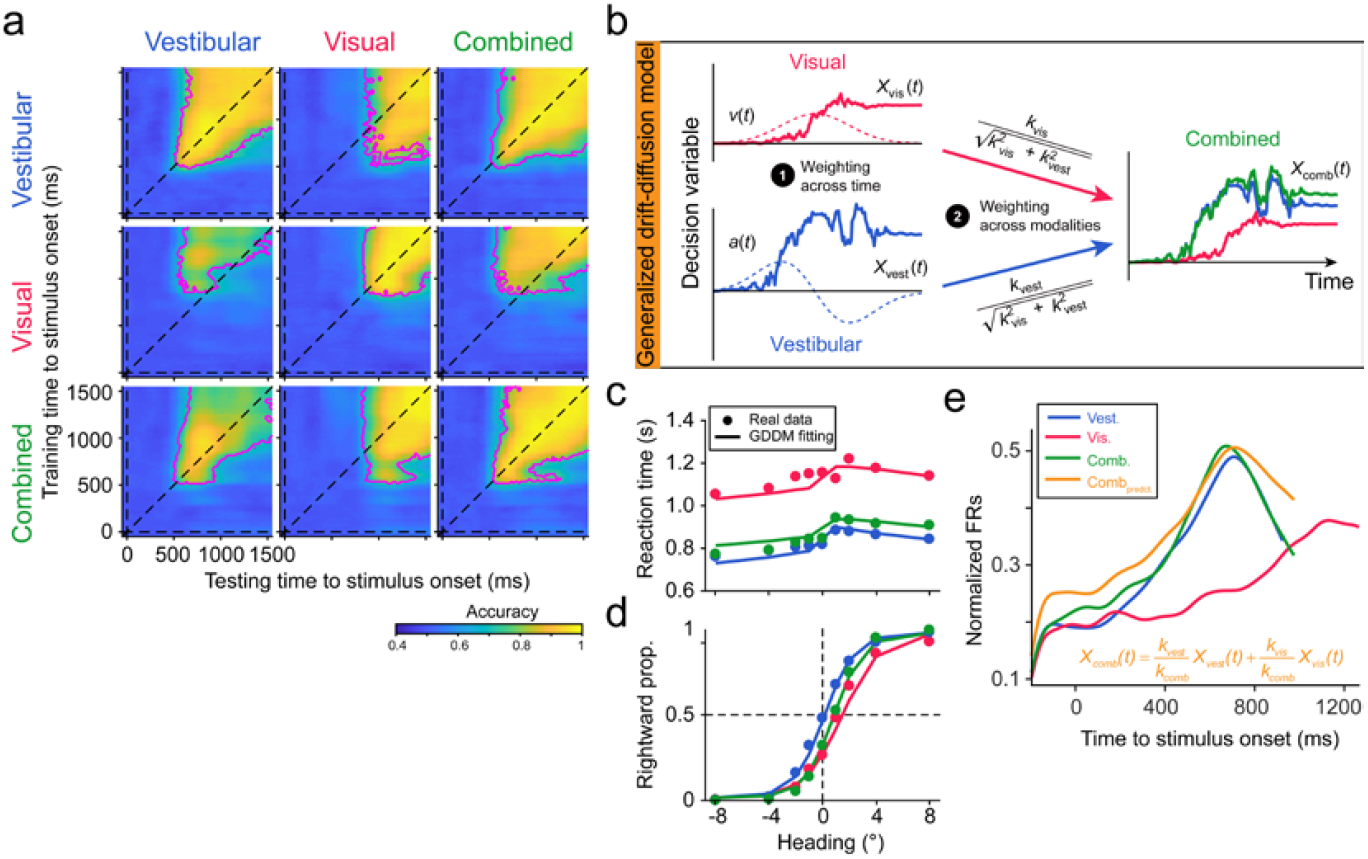
CN activity reflects animal multisensory decision behavior. **a**, Decoding accuracy of cross-temporal Lasso classifiers presented as heat maps. Subpanels on the principal diagonal are within-modality decoding, while those off the principal diagonal represent across-modality predictions. For each subpanel, the ordinate represents center of the time window used to train decoders, while the abscissa marks the testing time. Magenta lines outline the areas with accuracy significantly higher than chance level (50%, p < 0.01, permutation test). **b**, Schematic of the GDDM. Plot modified with permission from Drugowitsch et al.^36^ **c**, Chronometric curves. Dots are measured data from monkey M. Lines are fitting curves from GDDM. **d**, Psychometric curves. Labels are same as **c**. **e**, Population firing rates of CN (normalized). Orange line represents predicted bimodal response from GDDM. Blue, red, and green lines are measured CN responses in vestibular-only, visual-only, and bimodal condition, respectively.

The second analysis was to more directly assess whether CN dynamics could show optimality as seen in the behavior. Previous research has proposed a remarkable framework of generalized drift-diffusion model (GDDM, Fig. 4b, see Method), which can successfully explain human subjects’ heading performance under reaction-time (RT) context when both factors of choice correctness and response time were taken into account to assess optimality^36^. To do so, we trained a third monkey (M) to perform a RT version of the same heading discrimination task, in which the only difference was that the animal was allowed to make saccadic choice at any time right after the stimulus onset, without having to wait for the disappearance of the fixation point indicating the go-signal. In such a situation, we found the animal showed different response time across modality conditions. In particular, response time was significantly longer in the visual condition (∼1150 ms on average), compared to that in vestibular (∼820 ms on average) and combined (∼850 ms on average) condition (Fig. 4c, dot symbols). These patterns were similar to the human study^36^. GDDM well fitted behaviors including both of the chronometric (Fig. 4c, solid curves) and psychometric curves (Fig. 4d, solid curves). Importantly, the fitted drift rate, k-values (see Method), representing subjective stimulus sensitivity, were close between bimodal condition and prediction from optimal integration theory (Table 1), suggesting that monkey M integrated cues near optimally under reaction-time context. After acquiring k-values, the predicted moment-to-moment bimodal response in CN was consequently calculated, which turned out to be very similar with the real neurophysiological data (Fig. 4e, dark yellow versus green curves), supporting that CN dynamics could reflect the animals’ behavioral performance within the GDDM framework under more complex context when multiple factors need to be considered including response time, multimodal cues, and performance accuracy.

**Table 1.**
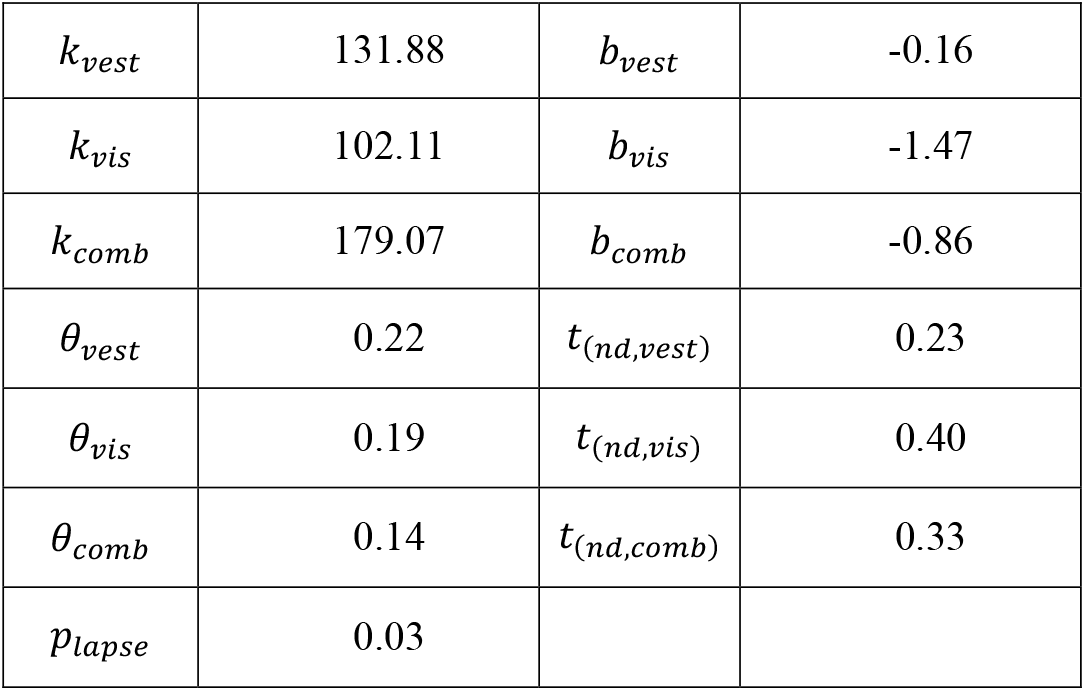
Fitting results of parameters in GDDM.

### Unilateral chemical inactivation in CN biased perceptual choice

We then examined causal-link roles of CN activity and the animal’s perceptual performance. Specifically, CN’s activities were manipulated through a number of methods, including chemical inactivation to test essential contributions, and electrical microstimulation to test sufficient contributions.

We first applied muscimol, agonist of GABA_A_ receptor to non-selectively suppress CN’s overall excitatory activity (Fig. 5a). We found that unilateral inactivation (monkey D, 5 cases; monkey F, 11 cases; Fig. 5b,c), but not saline control (monkey D, 6 cases; monkey F, 9 cases; Fig. 5f) induced significant ipsilateral bias in both animals’ choice. This ipsilateral bias was concordant with our recording data showing that majority of CN neurons preferred contralateral choice (Fig. 2a). In contrast, unilateral muscimol inactivation in LIP and FEF surprisingly produced negligible effect (monkey D, 3 cases; monkey F, 7 cases; Fig. 5d,e), although neurons in these two areas also mainly preferred contralateral choice (Extended Data Fig. 2b). Other than PSE, muscimol application did not significantly affected perceptual sensitivity (Extended Data Fig. 6a), or history information usage strategy (Extended Data Fig. 6b,c).

**Fig. 5.**
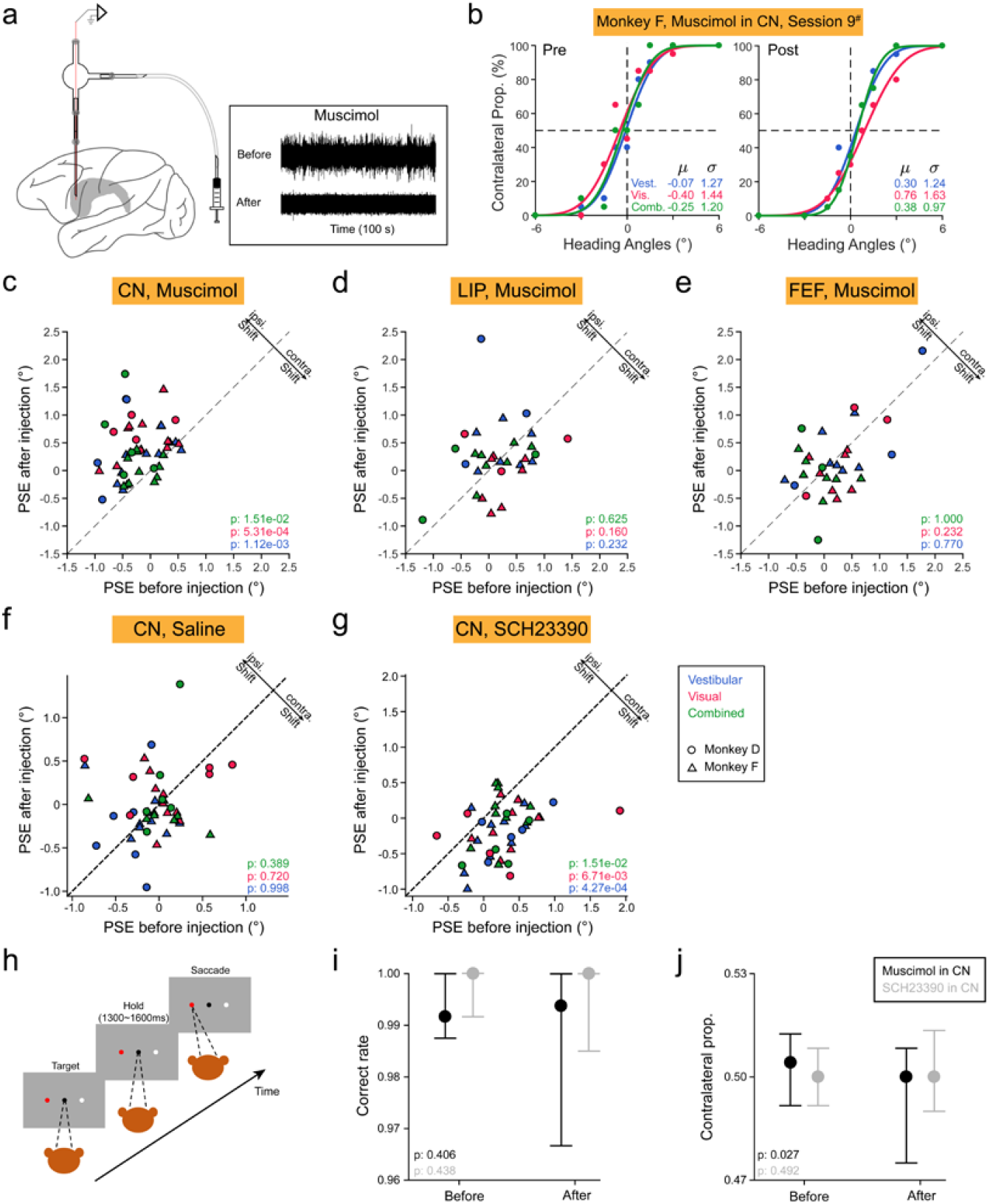
Unilateral inactivation of CN biases animal choice. **a**, Chemical injection into targeted areas including CN, LIP and FEF. Inset shows an example case of diminished neural activity after muscimol delivery. **b**, Example of one inactivation experiment, showing performance before (left plot) and after (right plot) muscimol unilateral injection in CN. **c-g**, Comparison of PSE before and after unilateral muscimol injections in CN (**c**), LIP (**d**), FEF (**e**), saline-injection control (**f**), and SCH23390 injections in CN (**g**). Shapes represent data from different monkeys. Colors are different modality conditions. All p values are from Wilcoxon signed-rank test; significance level, 0.05. **h-j**, Color guided saccade task (**h**). Comparison of accuracy (**i**) and PSE (**j**) before and after muscimol (black symbol) or SCH23390 (gray symbol) injection. Dot and error bar indicate median and 95% confidence interval. P-values are from Wilcoxon signed-rank test; significance level, 0.05.

Because CN is strongly innervated by midbrain dopaminergic inputs^37^, we wonder how selectively modulating dopamine receptors of CN neurons may affect the animals’ task performance. Thus, we then applied antagonist of D1 receptor, SCH23390, unilaterally into CN. We found that this manipulation induced significantly more contralateral choice (monkey D, 5 cases; monkey F, 10 cases; Fig. 5g), opposite to muscimol-mediated bias direction, implying that dopaminergic inputs into CN could modulate evidence accumulation process. Finally, similar to muscimol, blocking D1 receptors neither impacted heading sensitivity (Extended Data Fig. 6d) nor trial-history usage strategy significantly (Extended Data Fig. 6e,f).

These chemical-inactivation experiments suggest that CN plays a critical role in sensory-motor decision task, yet one possibility is that the observed causal effects may be mainly due to motor-report instead of cognitive perception, particularly when considering that CN is well-known for modulating movement executing^12, 38^. We think this is less likely to be the case in our study based on a few observations. First, chemical inactivation mainly affected difficult trials when heading directions were around the reference, which required more cognitive load compared to easy trials with large heading directions (Fig. 5b). Second, we further trained the animals to perform a simple color-selection task that required much reduced cognitive load than the fine heading discrimination task (Fig. 5h). In this controlled task, neither muscimol nor SCH23390 significantly influenced saccadic behavior any more (Fig. 5i,j), supporting that perturbation of CN activity mainly affected the animals’ cognitive process.

### Electrical microstimulation in CN indicated sufficient role

The third method for causal-link test was to use electrical microstimulation (50∼80 μA, 300 Hz, cathode-leading biphasic, Fig. 6a). Microstimulation was typically thought to identify sufficient contributions by artificially activating neural activity^39, 40^, although it still remains a debate about its exact impact on evoked neural dynamics^41, 42^. Specifically, in each experimental session, microstimulation was applied unilaterally in CN during the stimulus duration period, randomly for half of the total trials. Significance of PSE and threshold change in the psychometric functions were assessed by bootstrap test (Fig. 6b,c). Overall, microstimulation in CN was able to significantly bias PSE in a large proportion of cases (monkey D: 38/70 = 54.3%; monkey F: 52/76 = 68.4%, Fig. 6d). This proportion could be underestimated because of adaption of the currents^43^ (Extended Data Fig. 7f). The effect was fairly consistent across sensory modality (Extended Data Fig. 7b). In contrast to PSE, microstimulation only modestly affected psychophysical threshold (Extended Data Fig. 7a), indicating that microstimulation in CN mainly introduced signals instead of noise^44^, supporting that CN sufficiently contributes to the animals’ perceptual choice.

**Fig. 6.**
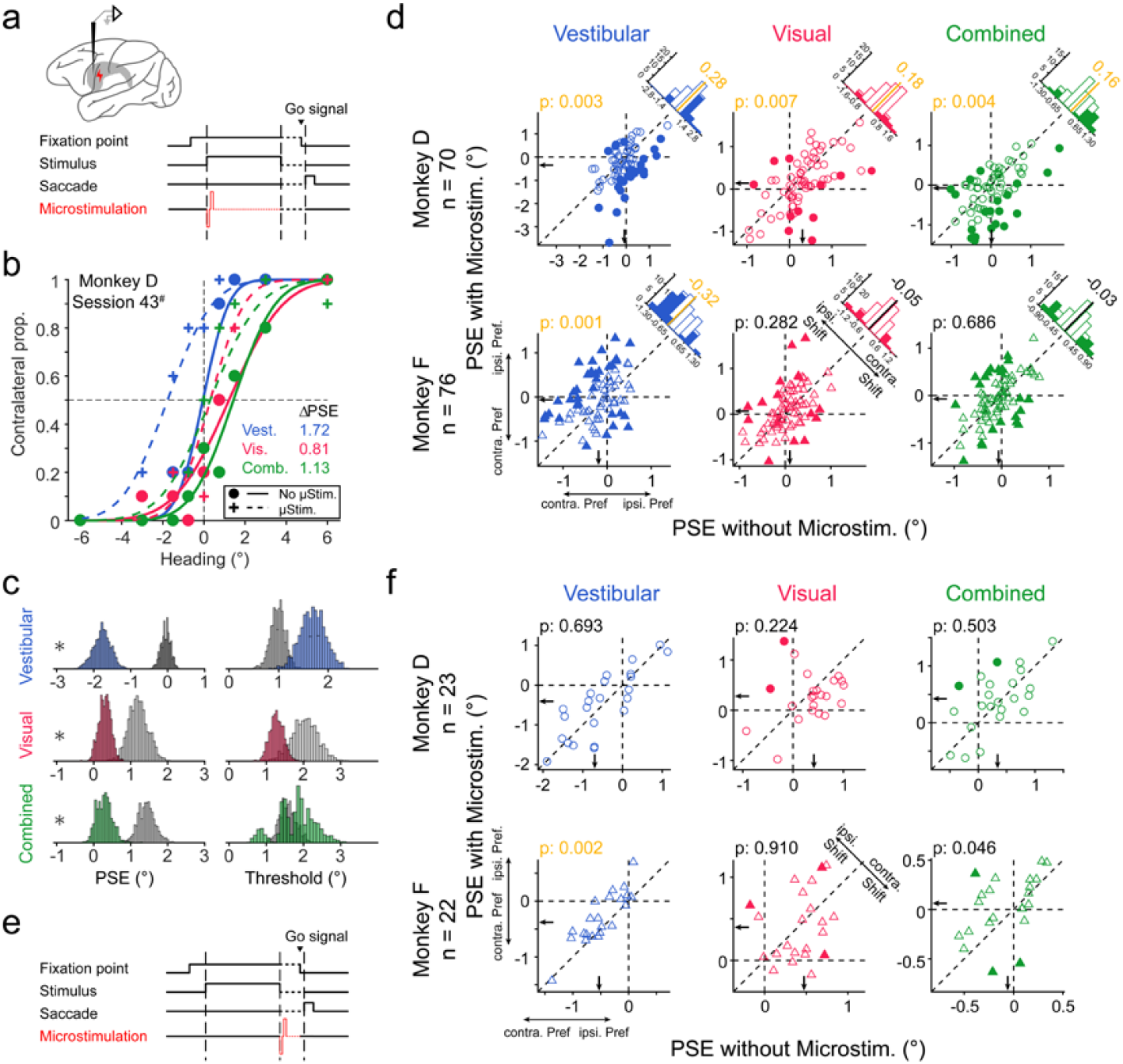
Unilateral microstimulation in CN affects animal choice. **a**, Schematic of unilateral microstimulation applied in CN during the stimulus-presentation period (1.5 s). **b**, Example of one experiment with microstimulation (dashed curves and cross symbols) and without microstimulation (solid curves and dots). **c**, Distributions of PSE (left column) and threshold (right column) constructed from bootstrap for microstimulation (color) and without microstimulation (gray) trials. Asterisk indicates significant change of the PSE or threshold (p < 0.01, bootstrap test). **d**, Comparison of PSE with (ordinate) and without (abscissa) microstimulation. Black arrows mark the median PSE. Marginal histogram counts frequency of ΔPSE. Solid bars indicate significant shift and open bars are insignificant change. **e-f**, Microstimulation applied during the delay period when stimulus is absent (**e**). Comparison of PSE with (ordinate) and without (abscissa) microstimulation (**f**).

We noticed, however, some inconsistence in PSE shift direction between the two animals. Specifically, microstimulation tended to induce more contralateral choice in monkey D (Fig. 6d, top row), but more ipsilateral choice in monkey F (Fig. 6d, bottom row). This individual difference could not be explained by factors such as anatomical location (Extended Data Fig. 7c,d), or coding properties of the stimulated sites (Extended Data Fig. 7e). To test other possibilities, we applied microstimulation during the delay period after stimulus offset and before saccadic choice (Fig. 6e). We found in such a case, microstimulation effects were also different between the two monkeys. In particular, while microstimulation effect was absent for monkey D, the effect remained for monkey F (ipsilateral bias, Fig. 6f). Thus, unlike the consistence of bias directions induced by chemical inactivation, microstimulation appeared to induce more confound results that might be due to different strategy of the animals, or limitations in the technique per se (see more in Discussion).

## Discussion

In the current study, we find neural manifolds in CN strikingly differ from that in LIP/FEF, although they exhibit similar low-dimensional task variables encoding pattern (Fig. 2). This cortico-subcortical distinction may be due to different single-modality inputs into individual areas (Fig. 3), but not to animal choice behavior (Extended Data Fig. 4). This finding challenges the general hypothesis that CN choice signals inherit and reflect sensory-motor association cortices. We then demonstrated that CN population activity is representing animal multisensory behavior in the task (Fig. 4), and that manipulating CN responses is able to bidirectionally affect monkeys’ decisions (Fig. 5, Fig. 6). Thus, our findings provide comprehensive evidence supporting that CN per se is of vital importance for perceptual decision-making function.

### Higher-dimensional paradigms help to dissect representational redundancy

It is common to see many brain functions are mediated by a distributed network involving multiple regions that share seemingly similar properties, leading to the “redundant” hypothesis^1-3^. One obvious benefit of this redundancy is that it helps to deal with dysfunction when some of the nodes within the network are hampered. Alternatively, the seemingly redundant nodes may actually carry distinct traits that can only be unraveled under more complex contexts. Here we provide clear evidence, showing that for perceptual decision-making, by adding an additional dimension of modality input, the subcortical area CN could become dramatically different from the frontal and parietal association cortices in low-dimension manifolds embedded in the higher-dimensional neural space. In particular, neural trajectory under bimodal condition is largely biased towards vestibular in CN, in contrast to a clear visual-bias in LIP and FEF. Aided by RNN simulations, our results suggest that on top of the role as a relay station for the cortico-striatal circuitry as in conventional view, CN could receive sensory evidence from elsewhere, for example, thalamus^45^, or other sensory cortices^46-48^, leading to subcortical-cortical heterogeneity. We expect that benefits the higher-dimensional paradigm bring in are not limited to perceptual decision-making studies. For example, future works can also parse the distributed representations in working memory, and other.

### Non-visual dominant CN may reflect animal decision strategy

Our perturbation experiments point out that the primate CN plays critical roles, both essential and sufficient, in the perceptual decision-making task, consistent with results from studies on rodents^25^. Considering that CN shows non-visual (vestibular) dominant neural state during bimodal condition when both single cues overall share similar cue-reliabilities, neither similar to the frontal (FEF) and parietal (LIP) sensory-motor cortex, nor to the polysensory area (MSTd), a question is what does this imply for CN in multisensory perception and decision-making? We think this finding is consistent with a few behaviors observed in previous studies.

First, we found CN population dynamics are consistent with prediction using parameters regressed from the generalized drift-diffusion model (GDDM). It’s noticeable that the fitted drift rate, reflecting speed of evidence accumulation, is higher for vestibular-only than visual-only condition, indicating an intrinsically higher sensitivity to vestibular stimulus^36^. Second, the vestibular bias may explain “vestibular-overweighting”, as reported previously in both human and non-human primate studies by using spatially conflict visuo-vestibular heading stimuli. Specifically, subjects’ PSE during bimodal condition with conflict unimodal cues is biased towards vestibular, even when the two unimodal cues carry matching reliability and analogous perceptual sensitivity (psychophysical threshold)^29,49^. Future experiments could be conducted by using cue-conflict task and examine CN’s role in the animals’ behavior. Third, the vestibular bias is consistent with the difference in response time under different modality conditions. Specifically, researchers have shown under a reaction-time version of task, human subjects tend to make choice faster in vestibular-only and bimodal stimulus conditions compared to that in the visual-only condition^36^. This is also exactly what we see in our current study by training one of the animals to perform the reaction-time task.

In addition to the vestibular bias in the neural state space, another feature in CN is that the Euclidean distance between different conditions ultimately decreases at the late stage of trials, unlike the sustained plateau in frontal and parietal cortices. This descending Euclidean distance actually also happens in the choice dimension, which may reflect overweighting of early phase stimulus and early termination of evidence accumulation during fixed duration task^50-52^. In our stimuli, visual sensory evidence becomes fairly weak at the late stage due to the Gaussian velocity motion profiles, and vestibular evidence is mainly concentrated at the early acceleration period. Thus, the later phase of sensory information may be largely neglected by the animals. Indeed, when switching to reaction-time context, the animal made saccadic choice much earlier (vestibular: ∼820 ms, visual: ∼1150 ms, bimodal: ∼850 ms) than the whole 1.5 second as required in the fixed-duration context. Thus, the dynamic choice signals in CN may be consistent with previous view that compared to cortex, CN is particularly important for decision formation at early stage^18^. In addition to choice signals, the descending modality information in CN may also be due to similar reasons: discrimination of modality (e.g. visual or vestibular) is not required in our task, therefore modality-related signals are ultimately faded at the end of trials. It would be interesting to test in future experiments by requiring subjects to discriminate modality conditions, in addition to heading stimuli, and examine whether related CN activity may sustain consequently.

### Dopaminergic modulation in CN is essential for perceptual decision

It has been recognized that dopaminergic regulation in neocortex is essential for many cognitive functions, such as working memory and associative learning, as indicated by selectively injecting chemical ligands of dopamine receptors into the prefrontal cortex^53-56^. Here, we showed that suppressing CN D1 receptors using antagonist consistently biased animals’ choice toward contralateral side, opposite to muscimol inactivation. Out result is thus concordant with the finding that low dose of D1 receptor antagonist typically elevates neuronal activity^57, 58^.

It is well known that the basal ganglia circuit contains two pathways of direct and indirect that is associated with D1 and D2 receptor, respectively. The two pathways are likely responsible for complimentary and opposite control of decision behavior^59^. It would be interesting to further test D1 and D2 receptor pathway in future experiments, as has been performed in rodents using state-of-art technique^13, 59^. For example, a recent work developed a recombinase-free retrograde adeno-associated virus (AAV) dependent strategy to precisely modulate CN direct MSNs^60^. It would be promising to identify primate CN circuit-specific contributions to cognitive functions in future studies.

### Implications of microstimulation experiment

Electrical microstimulation has been widely used in primates to identify causal roles of neural activity in perceptual choice^23,61, 62^. The technique is quite successful when applied in sensory areas, as evident by induced PSE shift in the direction consistent with the labelled-line of the stimulated neurons (encoded feature)^44^. The other advantage of applying microstimulation in these sensory areas is that in primates, neuronal signals sharing similar functions are typically well clustered in a local spatial domain (e.g. a few microns)^44^. Microstimulation in higher-level areas, or less sensory-dominant areas could be trickier, because different neuronal signals could be more mixed together, and they are less spatially clustered^23^. Indeed, in the current study we found although microstimulation in CN frequently induced significant PSE shift, confirming its sufficient causal role, the bias direction was not consistent between the monkeys. In particular, the ipsilateral bias is counterintuitive, based on the fact that majority of CN neurons prefer contralateral choice. This result, however, may be consistent with previous studies: CN may contain two components that show opposite preference for perceptual and motor process ^23, 63^. Other factors may also confound microstimulation result. For example, the electrical currents are easily spread, and subsequently activate passing fibers, leading to nonlocal effects ^41, 42^. Furthermore, electrical currents lack specific targeting, and may impact direct and indirect pathway differently, consequently generating heterogeneous result. Optogenetic or chemogenetic tools, that are typically much more relieved from these issues, may be applied in future experiments to further test sufficient roles of certain CN populations in perceptual decision tasks.

## Acknowledgements

We thank Wenyao Chen for monkey care and training, and Ying Liu for C++ software programming. We thank Jan Drugowitsch for kindly offering GDDM code. This work was supported by grants from the National Science and Technology Innovation 2030 Major Program (2022ZD0205000) to Y.G.

## Author Contributions

Z.Z., H.H. and Y.G. conceived the project and designed the experiments. Z.Z. and Y.X. performed the experiments. C.Z. and Z.Z. constructed RNN simulations. Z.Z. analyzed the data. Z.Z. and Y.G. wrote the manuscript.

## Declaration of interests

The authors declare no competing interests.

## Extended data

Extended Data Fig. 1-7.

## Methods

### Animals and surgeries

Three adult monkeys (*Macaca mulatta*), monkey D, F and M, weighting 7∼10 kg, were trained in this work. Details about animal surgery and training procedures have been described previously^26^. In brief, monkeys were chronically implanted with a 6-cm lightweight plastic ring, which was served as recording chamber as well as for head restraint. Then, monkeys sat comfortably in a customized primate chair fixed within a visuo-vestibular virtual reality system. All animal procedures were approved and supervised by the Animal Care Committee of Center for Excellence Brain Science and Intelligent Technology, Chinese Academy of Sciences.

### Apparatus

The visuo-vestibular virtual reality system consists of a motion platform (MOOG MB-E-6DOF/12/1000KG) and a LED display (∼30 cm of view distance and ∼90° × 90° of visual angle; Samsung ED55C) vertically mounted on the platform, to offer vestibular and visual self-motion stimuli, respectively. The stimuli were controlled by a customized C++ software and synchronized with electrophysiological recording (AlphaOmega SnR, Israel) and eye tracking (SR Research, EyeLink 1000 Plus, Canada) systems by TEMPO (Reflective Computing, U.S.A).

### Behavioral tasks

#### Memory-guided saccade task (MGS)

Similar to previous works^64^, monkeys were trained to perform a memory-guided saccade task to identify the response field (RF) of the recorded neuron. However, note that we didn’t use this task to screen neurons in the later heading discrimination task, considering that there is only weak correlation between strength of choice signal and spatial selectivity^10^. In brief, monkeys were required to fixate on a central fixation point for 100 ms to initiate a trial, after which a peripheral target flashed for 500 ms in one of the eight positions with 10° away from the display center. Monkeys were required to keep central fixation for another 1000 ms, before the fixation point disappeared, indicating a go signal for the memorized peripheral target.

#### Multisensory heading discrimination tasks

Monkeys initiated a trial by fixating on the central fixation point, after which two choice targets appeared symmetrically on each side of the display with an eccentricity of 10°. After a random delay (100∼300 ms), a 1.5-second, linear forward self-motion stimulus, defined by either visual, vestibular, or bimodal cues was provided in the horizontal plane with a small deviation from the reference of straight forward (0° heading). After the stimulus offset, the animals waited for another random delay (300∼600 ms), before the fixation point disappeared as a go-signal. The animals were then allowed to saccade toward one of the two choice targets to report whether their experienced self-motion direction (heading) was either in the leftward or rightward category. Correct answer led to reward of a drop of juice. For dead ahead trials (0° heading), reward was randomly delivered.

In each experimental block, there were three modality conditions: (1) vestibular-only, where self-motion stimuli were solely offered by physical motion of the motion platform; (2) visual-only, where self-motion stimuli were simulated via optic flow on the display, while the motion platform was stationary; (3) cue-combined, where congruent vestibular and visual cues were provided synchronously. Similar to previous works^27, 65^, all stimuli followed a Gaussian velocity with a biphasic acceleration profile to simulate transient self-motion in the environment. Note that to best see a cue combined effect, we have downgraded the visual cue reliability through coherence^66^ (10%-30%) to match the vestibular, so that psychophysical threshold under the two single cue conditions would be roughly similar^26^. Task difficulty was determined by heading angles (monkey D: ± 8°, ± 4°, ± 2°, ± 1°, 0°; monkey F: ± 6°, ± 3°, ± 1.5°, ± 0.75°, 0°; monkey M: ± 8°, ± 4°, ± 2°, ± 1°, 0°). Thus, each repetition contained 27 interleaved trials (3 stimulus modality × 9 heading angle), and each repetition was typically repeated for more than 20 times, leading to more than 500 trials.

Other than the fixed-duration task (1.5 second), a reaction-time version of the task was also introduced for monkey M. Specifically, the animal was allowed to make choice before the go-signal appeared, without having to maintain central fixation across the whole stimulus duration period (1.5 second).

#### Color selection task

The aim of this task is to test motor execution and bias. The animals were only required to saccade to the red target to earn a liquid reward. The structure of the trial was identical to the heading discrimination task, but with no self-motion stimulus appearing.

### Electrophysiology

#### Caudate mapping

Area mapping was made through cross-validation between structure MRI and physiological properties. Along each electrode penetration, baseline changes of electrophysiological signals were carefully monitored to determine transition patterns of gray/white matter. As illustrated by Extended Data Fig. 1g, the recording sites mainly cover head and body parts of CN (AC: -12∼8 mm), similar to previous works^10, 67^.

#### On-line recording and Off-line sorting

Once encountering a single neuron in CN, we identified its response field using MGS task. Three epochs were focused: (1) visual response period, 75∼400 ms after appearance of the peripheral visual target; (2) memory period, 25∼900 ms after the peripheral target disappeared; (3) pre-saccade period, 300∼50 ms before saccade initiation. If this neuron showed significant spatial selectivity during any of the three windows, we then used it to guide placing the two choice targets during the following heading discrimination task (IN and OUT). If a neuron did not show significant modulation during the MGS task, we also ran heading discrimination task, albeit with the choice targets placed in the horizontal meridian. All online recorded data were performed off-line sorting, using Spike2 for single-channel data and Kilosort-2.0 for linear-array data.

#### Selection of putative medium spiny neuron

Three subtypes, including the median spiny neuron (MSN), tonically active neuron (TAN) and fast-spiking neuron (FSN), account for more than 95% of CN neurons^68^. Among them, MSNs are the projection neurons that we aim for, yet they are not easily targeted especially for primate electrophysiology. Thus, we tried to identify MSNs from TANs and FSNs according to a number of indicators^30^. (1) Waveform. Compared to MSNs, FSNs tend to have narrower spike waveforms, measured by the shorter peak-to-trough time. (2) Spontaneous activities. While spontaneous activity of MSNs and TANs are often as low as less than 10 Hz, FSNs usually reach a very high level. We calculated the spontaneous firing rates for each neuron using mean activities of 600∼800 ms after reward feedback in the main decision-making task. (3) Firing patterns. While TANs usually fire tonically, MSNs and FSNs often present phasic firing. We measured this through distributions of the inter-spike interval, calculating *In*(*ISI*_*median*_/*ISI*_*mean*_).

The single-channel data were recorded using two systems, AlphaOmega SnR and CED, and outlier detection was conducted separately for each system to avoid system difference (Extended Data Fig. 1a). Note that inclusion of all recorded neurons didn’t change our conclusions, because MSNs occupies majority of the CN neurons.

### Reversible chemical inactivation

Two drugs, Muscimol·HBr and SCH23390·HCl (from Sigma-Aldrich), were first dissolved in saline to the concentration of 10 mg/mL and 0.25 mg/mL, respectively. They were accurately injected (∼4 μL) into the targeted areas at a speed of 0.15 μL/minute using the hand-made injectrode with an electrode embedded. Because the half-life eliminations of the two drugs are very different, their effects on the animal behavior can last differently, which is about 12 hours for muscimol, and less than one hour for SCH23390. Thus, we applied two consecutive injections with interval of 48 hours for muscimol experiments, and 24 hours for SCH23390, respectively.

### Electrical microstimulation

For a traditional microstimulation experiment particularly applied in sensory cortices, it’s necessary to find the neuronal cluster with similar electrophysiological properties near the electrode tip^61, 62, 69^. However, it has been reported that there is no such cluster in CN^10^. So we did not systematically measure clustering to guide microstimulation, similar to previous work^23^. For the sake of comparison with others’ work^23, 63^, the current was set to be negative-leading bipolar pulses, 300 Hz, 50∼80 μA. These parameters would not evoke saccades^23, 70^. In a microstimulation block, 50% trials were randomly chosen to deliver the electric current, making it unlikely for the animals to quickly adapt strategy.

### Data analysis

#### Psychophysics

To quantitatively measure the animals’ behavioral performance in the perceptual heading discrimination task, we constructed psychometric curves by plotting the proportion of “rightward” choice as a function of heading angles. These curves were fit with cumulative Gaussian functions, whose mean (μ) and standard deviation (σ) were defined as point of subjective equality (PSE) and psychometric threshold, respectively. The Bayesian optimal prediction of psychometric threshold in the bimodal condition for each session was calculated by

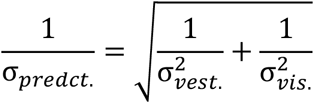

where σ_*vest*._ and σ_*vis*._ indicate psychometric thresholds measured in vestibular-only and visual-only conditions, respectively^26^.

#### Choice and modality preference

To construct peri-stimulus time histogram (PSTH), raw spike trains were aligned to events of stimulus onset, saccade onset, and feedback onset. Firing rates were computed in every 10-ms time window and smoothed with a Gaussian kernel (σ = 50 ms). Next, all correct trials were grouped according to animals’ choice (IN versus OUT target). When averaging, PSTHs were normalized for each cell across time and modalities by

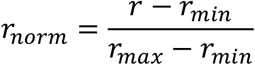

where *r*_*max*_ and *r*_*min*_ were respectively maximum and minimum firing rates for each neuron. To quantify strength of choice signals, we used receiver operator characteristic (ROC) analysis to define an index of choice divergence (CDiv)^27^. CDiv ranged from - 1 to 1, with 1 meaning always stronger firing when IN target is chosen, and vice versa for -1. CDiv = 0 indicates no choice preference. Grand CDiv was also computed by using firing rates across the whole stimulus duration to identify choice preference for each neuron in each modality condition.

Similarly, comparing firing rates between vestibular-only and visual-only condition, we defined modality preference (MDiv), with 1 indicating strong visual preference, and -1 indicating strong vestibular preference.

#### Direction discriminative index (DDI)

To estimate the strength of neuronal selectivity to directions in MGS task and passive heading task, DDI^27^ was computed as:

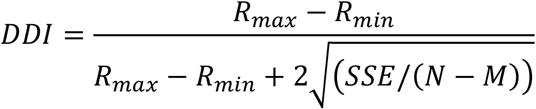

where *R*_*max*_ and *R*_*min*_ respectively indicate mean firing rates on the direction with strongest and weakest activities; *SSE* is the sum-squared error around mean responses; *N* is the total number of observations (trials), and *M* is the number of stimulus directions. Value of DDI ranges from 0 to 1, with the larger value indicating stronger selectivity.

#### Dimensional reduction of neural activities

To better visualize neural trajectories in low-dimensional subspace and demix task variables, we used three dimensionality reduction methods: principal component analysis (PCA), targeted dimensionality reduction (TDR), and demixed principal component analysis (dPCA).

For PCA, we first constructed a matrix ***r*** of size *N*_*unit*_ × (*N*_*T*modalities*2choices*_), whose columns contained firing rates of all single neurons during a specific time bin in each stimulus condition when one of targets was chosen. Then, standard singular value decomposition (SVD) was conducted to get left singular vectors, ***U***. The first three columns of ***U*** (3 PCs) were chosen to span a denoising subspace, onto which the population activities ***r*** were projected. In this three-dimension space, six neural trajectories (3 modalities and 2 choices) were presented as a function of elapsed time.

For TDR, a subspace of ***D*** was constructed using the first 15 PCs of PCA. Linear regression was then used to capture task-related variables (heading, choice, modality) in the neural state space. Specifically, z-scored firing rates of neuron *i* at time *t* were predicted as a linear combination of multiple variables:

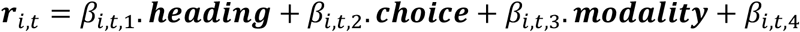

where ***r***_*i,t*_ is the z-scored firing rates of neuron *i* at time *t* in all trials, ***heading*** represented heading directions of all trial, ***choice*** included animal decisions (−1: OUT choice; 1: IN choice), and ***modality*** contained sensory modalities (1: vestibular-only; -1: visual-only). The regression coefficients *β*_*i,t*_ measured how much the trial-by-trial firing rates of neuron *i* at time *t* were dependent on the corresponding variables. After that, we denoised ***β***_***t***_ (includes *β*_*i,t*_ from all neurons) by projecting it onto denoising subspace *D*, and acquired 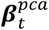. For each task variable, we then determined the time *t*_*v*_ when 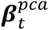 had the largest norm, so that the de-noised regression subspace ***β***_***v***_ was captured by that variable. Next, we orthogonalized each ***β***_***v***_ to get the task-related subspace through QR-decomposition. Last, the averaged population activities were projected onto this targeted subspace. More detailed descriptions can be found in the supplementary part of a previous work^35^.

For dPCA, we utilized toolbox published previously^32^. Briefly, we first built the neural data matrix *r* of size *N*_*unit*_ × *N*_*modality*_ × *N*_*choice*_ × *T* × *N*_*rep*._), where *N*_*unit*_, *N*_*modality*_, *N*_*choice*_ and *N*_*rep*._ respectively represent number of neurons, modality conditions, choices and repetitions. Then, decomposing the neural activities into five parts: condition-independent, modality-dependent, decision-dependent, modality-decision interaction-dependent, and noise through marginalization. The principal components relative to corresponding parts were found by minimizing the difference between the marginalized data and the reconstructed full data.

#### Cross-temporal Lasso decoders

To decode monkeys’ choice from CN population activity, a series of linear classifiers were trained and used as a function of time over the trial:

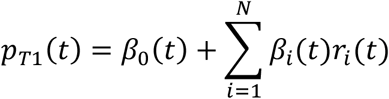

where *p*_*T*1_(*t*) represents the probability of choosing T1 target (IN choice) at time *t, r*_*v*_ indicates firing rates of neuron *i* at time *t, N* is number of neurons. Assuming sparse encoding, LASSO regularization was applied to prevent over-fitting when fitting each set of parameters ***β***.

Trials of each neuron were first categorized into 6 groups according to choice and modality, and firing rates in each trial in a sliding 100-ms bin (step size was 20 ms) was computed. 80 trials within each group were randomly selected for training, and the rest trial (>10) were used for testing. During training, the best regularization parameter of the model was determined by a 10-fold cross-validation. During testing, the model predicted T1 (IN) choice if the *p*_*T*1_(*t*) > 0.5 and T2 (OUT) choice if *p*_*T*1_(*t*) < 0.5 at any moment. This step was repeated 1000 times to assess significance over chance level.

#### Generalized drift-diffusion model (GDDM)

To predict firing dynamics in bimodal condition in reaction-time task, we used GDDM proposed previously^36^. In brief, the model assumes that both sensory and evidence accumulation processes are modulated by cue reliability:

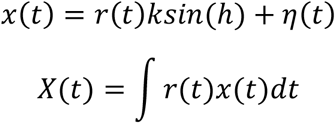

where *x*(*t*) and *X*(*t*) are respectively sensory evidence and decision variable at time *t* ; *r*(*t*) indicates moment-to-moment reliabilities of sensory stimuli; *k* denotes constant drift rate, measuring the subjective sensitivity to stimuli; *kin*(*h*) is the horizontal projection of heading; *η*(*t*) represents noise. Based on previous findings, the models assumes that subjects rely on vestibular acceleration but visual velocity to perform heading discrimination task^36^. Cue-reliability during bimodal condition was formulated as:

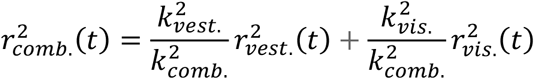

After well-trained, the optimal subjective sensitivity was given by:

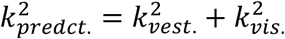

The accumulated evidence under bimodal condition could also be predicted from:

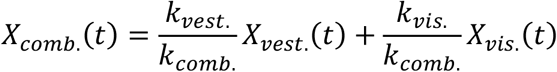

The model encompassed 13 parameters [*k*_*vis*._, *k*_*vest*._, *k*_*comb*._, *θ*_*vis*._, *θ*_*vest*._, *θ*_*comb*._, *b*_*vis*._, *b*_*vest*._, *b*_*comb*._, *t*_(*nd,vis*.)_, *t*_(*nd,vest*.)_, *t*_(*nd,comb*.)_, *p*_*lapse*_ ] that were used to fit chronometric and psychometric curves of all three modality conditions simultaneously. Among the parameters, *k* denotes drift rate; *θ* indicates decision threshold; *b* is heading bias; *t*_*nd*_ represents non-decision time; *p*_*lapse*_ means lapse rate.

#### Normalized shifts of point of subjective equity

To compare the effects of microstimulation across modal conditions, we normalized the change of PSE (Δ*PSE*) in each session by dividing the psychometric threshold under the condition without microstimulation. This generated Δ*PSE*_*norm*_, which was unitless.

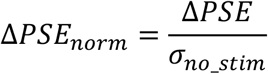

### Recurrent neural network simulations

We trained randomly initiated recurrent neural networks (RNN), comprising 256 nonlinear hidden units, a multisensory heading discrimination task analogous to that performed by monkeys. For all units, a tanh activation function is used:

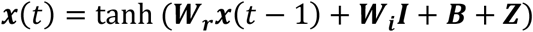

where matrices ***W***_*r*_ and ***W***_***h***_ denote recurrent and input weights, respectively; ***B*** represents activity offset; ***Z*** indicates that all units are affected by independently and identically distributed (i.i.d.) white noise, with a standard deviation of 0.1.

#### Network input and output

The network receives visual velocity but vestibular acceleration input simulating self-motion in 9 heading directions ([±8, ±6, ±4, ± 2, 0]). The stimuli last 1.5 s with a temporal resolution of 0.05-s. The sensory evidence is influenced by i.i.d. noise, with variance proportional to the input magnitude, leading to analogous signal-to-noise ratio between the two single cues, that is, visual and vestibular condition. Indeed, after training, the network produced analogous performance (psychophysical threshold) in visual and vestibular condition.

An output neuron reads out RNN’s population activity using linear weighted sum, and generates a binary choice at the end of each trial:

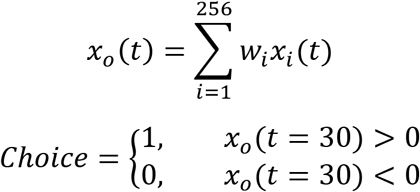

where *x*_*c*_ represents activity of the output neuron, and *x*_*v*_ indicates activity of the *i*th neuron in the RNN.

#### Network training

The weights of inputs-RNN, RNN-units, RNN-output are initialized through He initialization and updated simultaneously using Adam Optimizer and the back-propagation through time (BPTT). Loss in each time step is calculated using BCELoss combined with a Sigmoid layer (BCEWithLogitsLoss function):

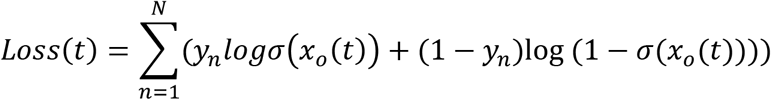

where *y*_*n*_ is the label of trial n; N is batch size; *σ* is sigmoid function.

## Extended data

**Extended Data Figure 1.**
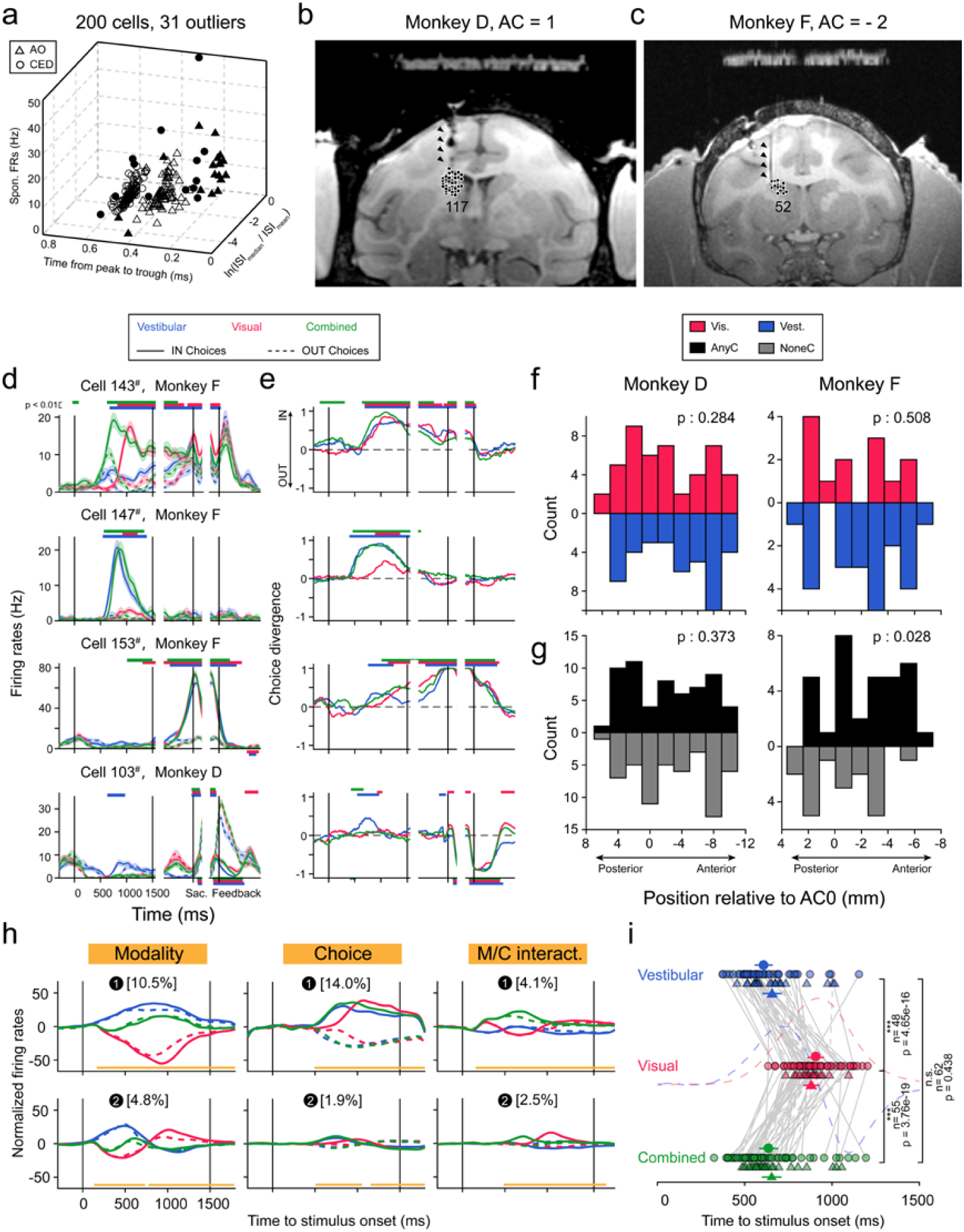
Anatomical distribution of CN MSNs and neuronal properties. **a**, Selecting MSNs using robust covariance estimate method. The 3-d coordinate indicates that electrophysiological properties are quantified through three metrics: x axis, time from peak to trough of spike waveform; y axis, spontaneous firing rates; z axis, median over mean of the inter-spike interval distribution. Shapes of the data points indicate recording systems (triangle: AlphaOmega SnR; Circle: CED); solid dots represent outliers detected by the algorithm. **b-c**, Representative coronal sections of monkey D (**b**) and monkey F (**c**). All recorded neurons were superimposed onto this section. Triangles mark the artifacts of electrodes from MRI scanning. **d**, PSTHs of four example neurons. For each cell, responses were respectively aligned to the stimulus onset (left subpanels), saccade onset (middle subpanels), and feedback onset (right subpanels). Blue: vestibular-only; Red: visual-only; Green: bimodal condition. Solid and dashed lines respectively represent trials where monkeys choose the IN and OUT targets with respect to the response field of the recorded neuron. Horizontal colored bars mark the time period when IN and OUT trials significantly differ in firing rates (p < 0.01, two-tailed t-test). Bar’s location indicates the relationship between their responses (IN > OUT, top; IN < OUT, bottom). **e**, CDivs of the same four neurons in **d**. **f**, Spatial locations for neurons with different modality preferences in the two animals. Only neurons with significant modality preference were included. Red and blue bars indicate visual and vestibular preferences, respectively. Abscissas represent the location relative to AC0 along the AP axis (positive, posterior to AC0; negative, anterior to AC0), and the ordinates count the frequency. All p values are from Mann-Whitney U-test. **g**, Spatial locations of neurons with and without significant choice preferences. All markers are same as **f**, except that colors here denote whether neurons have significant choice preferences (black, significant; gray, not significant). **h**, De-mixing task variables from CN population through dPCA analysis. Columns and rows respectively represent task-related parameters (modality, choice, and modality-choice interaction) and their dPCs (sorted by the proportion of explained total variances, ETV). For each subpanel, the percentage above means proportion of ETV; orange horizontal bars mark the period when corresponding variable can be significantly decoded from that dPC. **i**, Across-modality comparison of divergence time. Data points with error bars denote mean and SEM; gray lines connect data from the same neurons; *** and n.s. respectively represent p < 1e-03 and no significance; two-tailed paired t-test; significance level is 0.01.

**Extended Data Figure 2.**
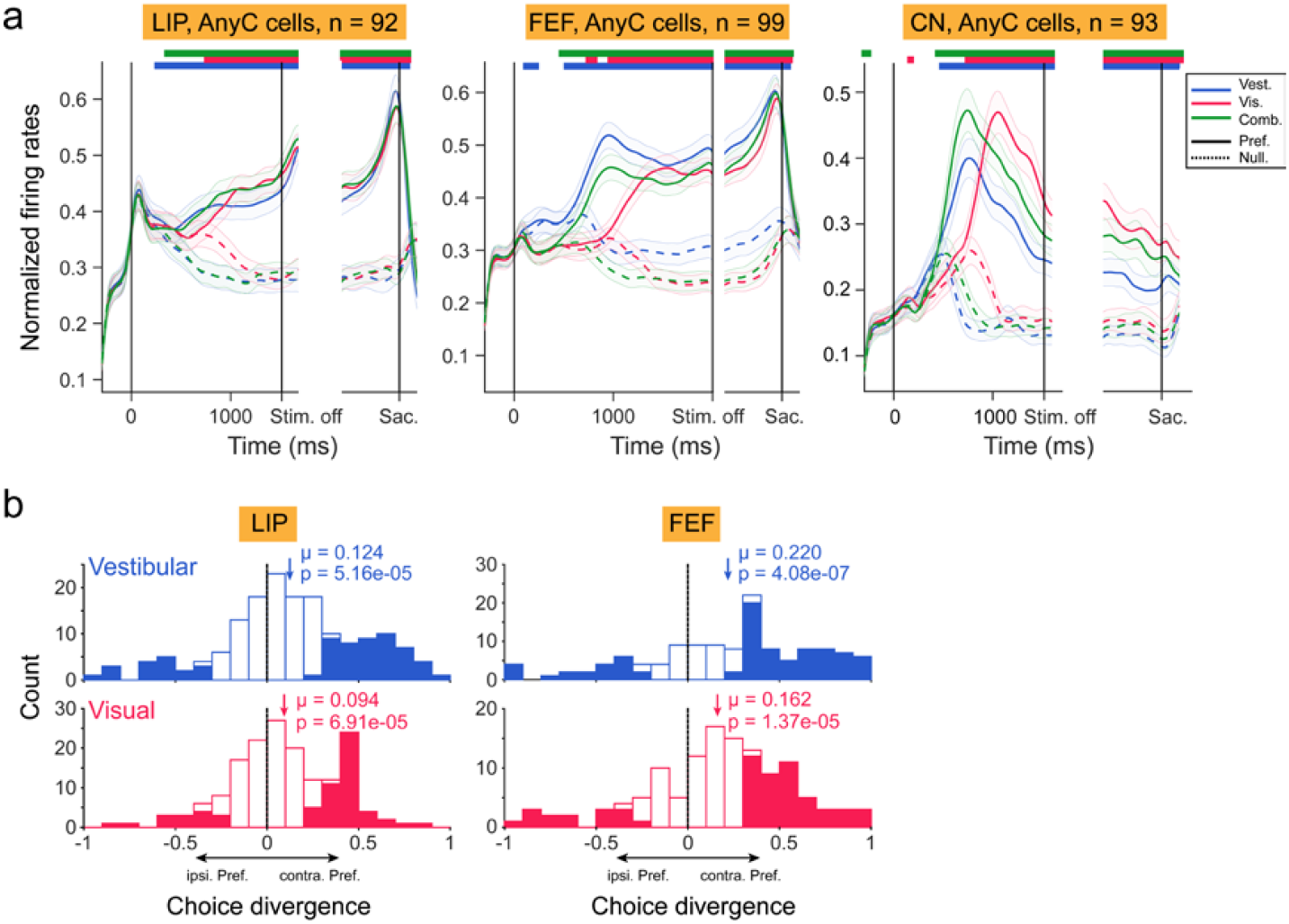
Choice related signals across areas. **a**, Population normalized PSTHs of decision-related neurons in LIP (left), FEF (middle), and CN (right). Only cells encoding choice signals in at least one modality conditions are included. Trials are sorted according to modality context and choice. Labels are same as Fig. 2b. **b**, Distributions of choice signals under single modality context in LIP (left column) and FEF (right column). Labels are same as Fig. 2a.

**Extended Data Figure 3.**
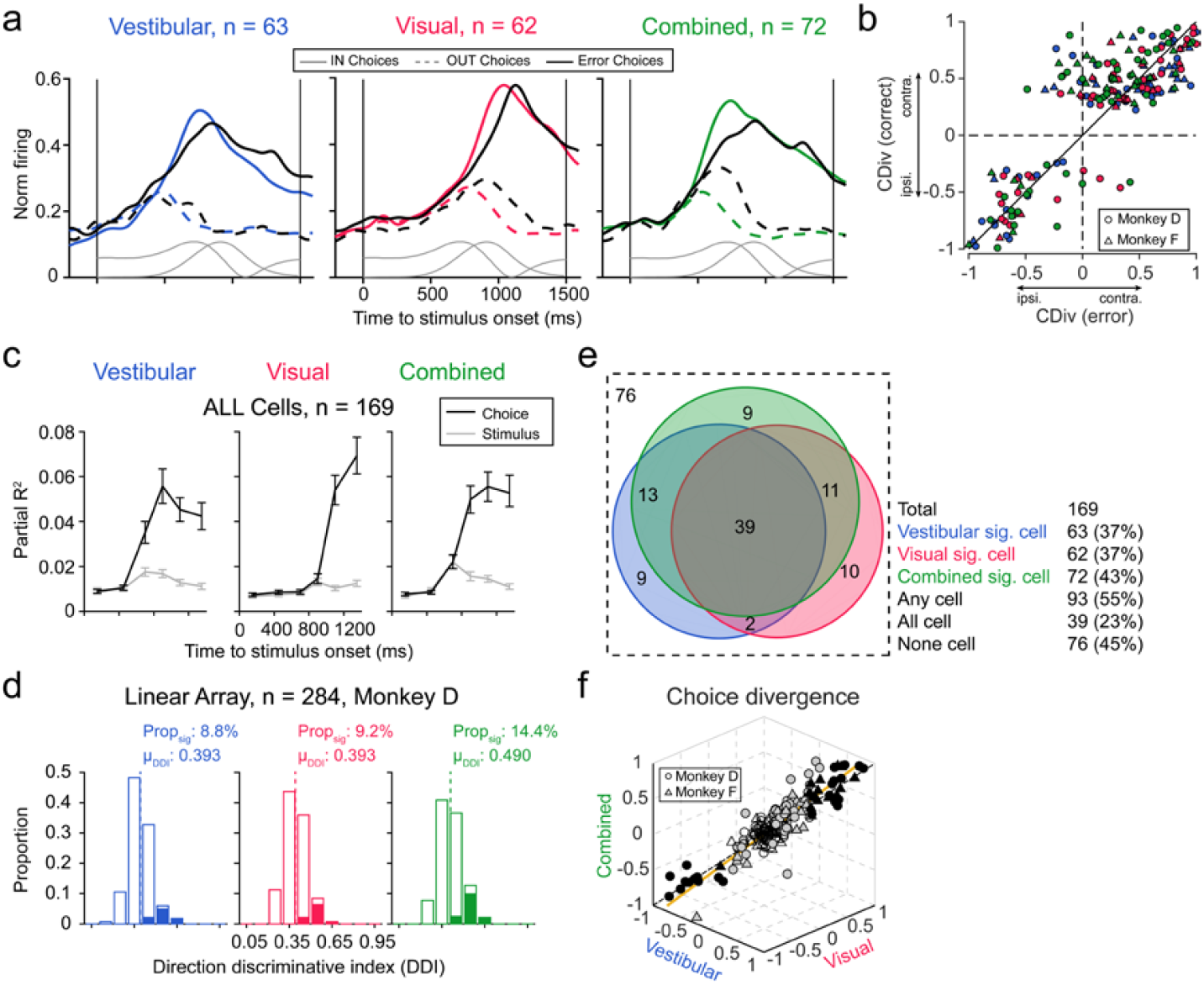
CN neurons are largely modulated by internal decision rather than external physical stimulus. **a**, Population average of normalized firing rates in correct (color curves) and error trials (black curves). Solid and dashed lines respectively denote trials where IN and OUT targets are chosen. Gray curves: motion stimulus profile. **b**, Comparing CDivs in correct versus error trials. Sign represents direction (positive: preferring contralateral choice; negative: preferring ipsilateral choice). Circles and triangles are data from monkey D and F, respectively. **c**, Partial correlation of neural responses with choice (black curves) and with heading (gray curves). Error bars represent SEM. **d**, Distributions of neuronal DDI measured during passive, fixation-only heading task. Solid bars indicate significantly tuned neurons (one-way ANOVA, p < 0.05). **e**, Classification of CN neurons according to grand CDiv. Number within each colored region marks the count of cells with significant grand CDiv under the corresponding combination of modal conditions. **f**, Correlation of CDivs across modality conditions. Circle and triangle dots are data from monkey D and F, respectively. Colors indicate how many modality conditions in which CDivs are significant (black, three; gray, one or two; empty, none). Orange line is the first principal component of PCA analysis; the first PC captures more than 88% variances; dashed black line is the diagonal.

**Extended Data Figure 4.**
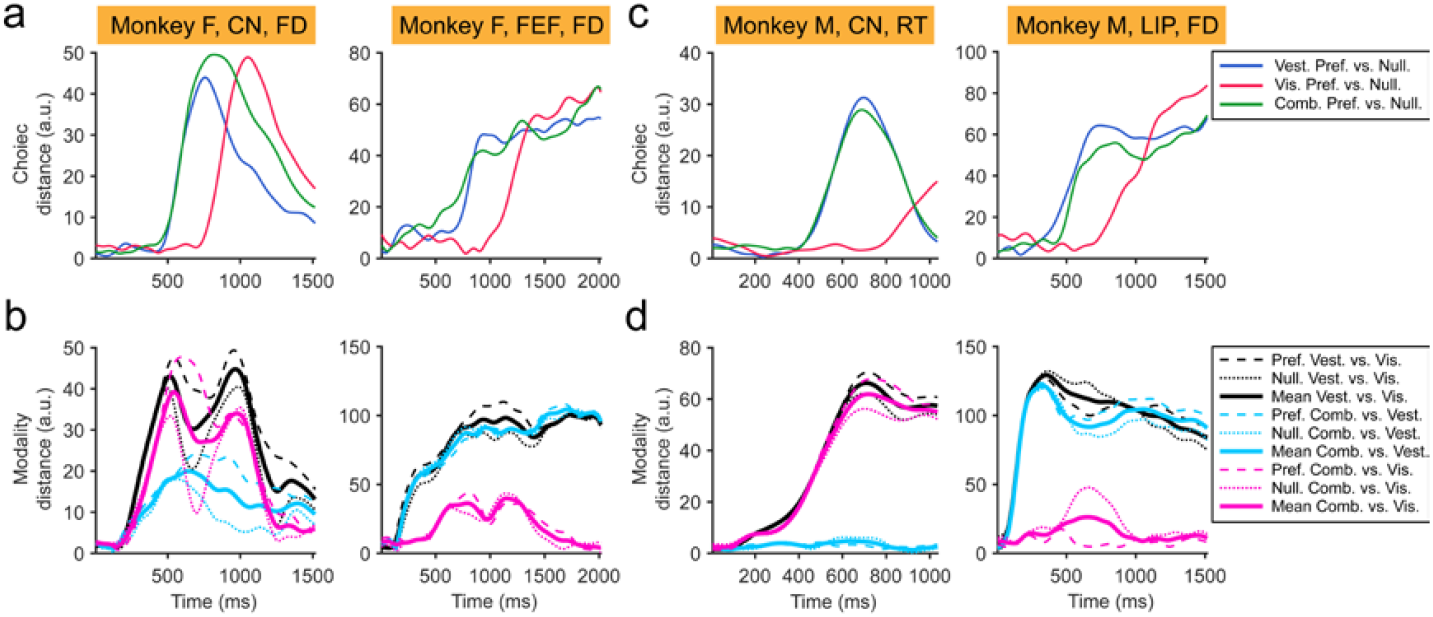
Comparing neural states across regions within the same monkey. **a-b**, Temporal dynamics of choice (**a**) and modality (**b**) distance in CN (left column) or FEF (right column) of monkey F performing the fixed-duration task. Labels are same as Fig. 3d. **c-d**, Temporal dynamics of choice (**c**) and modality (**d**) distance in (left column) CN of monkey M performing the reaction-time task, or in (right column) LIP of the same monkey but performing the fixed-duration task.

**Extended Data Figure 5.**
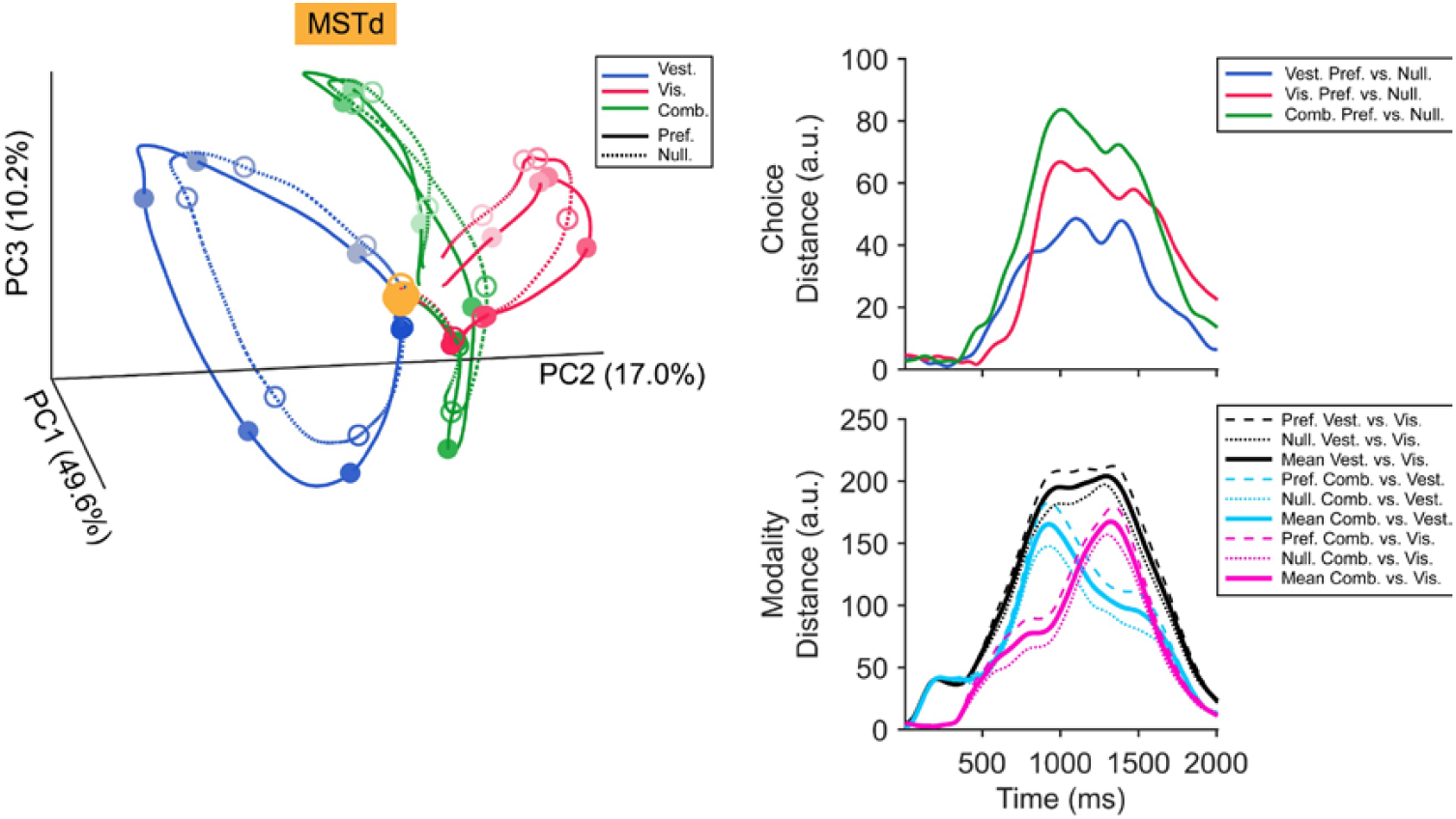
Neural state trajectories of area MSTd. Labels are same as Fig. 2.

**Extended Data Figure 6.**
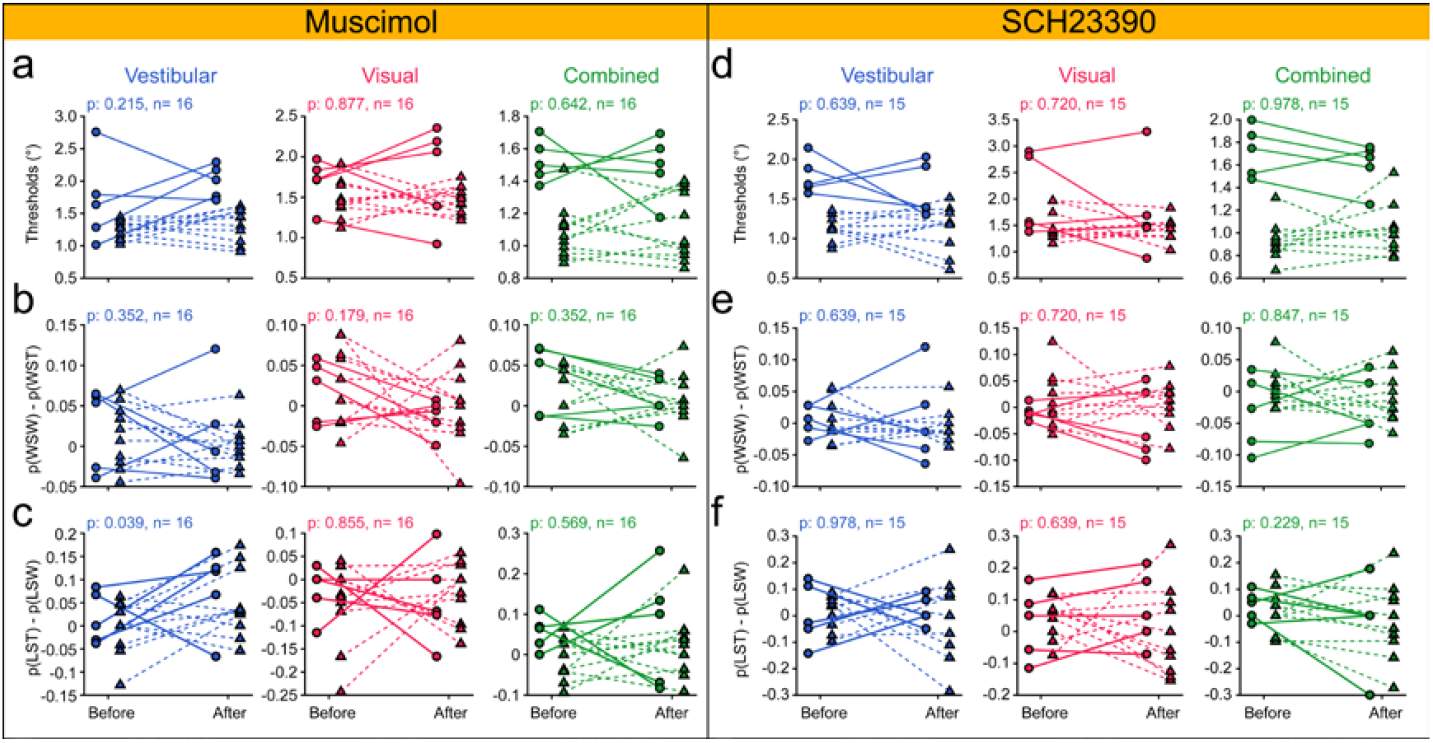
Effects of unilateral chemical inactivation in CN on perceptual sensitivity and history strategy. **a**, Psychometric thresholds before and after injecting muscimol. Different symbols represent data from the two animals. P-value is from Wilcoxon signed-rank test; significance level is 0.05. **b**, Win-switch versus win-stay proportion before and after injecting muscimol. Labels are same as **a**. **c**, Lose-stay versus lose-switch proportion before and after injecting muscimol. Labels are same as **a**. **d-f**, Same format as in (**a-c**), but for D1-receptor antagonist SCH23390 injection in CN.

**Extended Data Figure 7.**
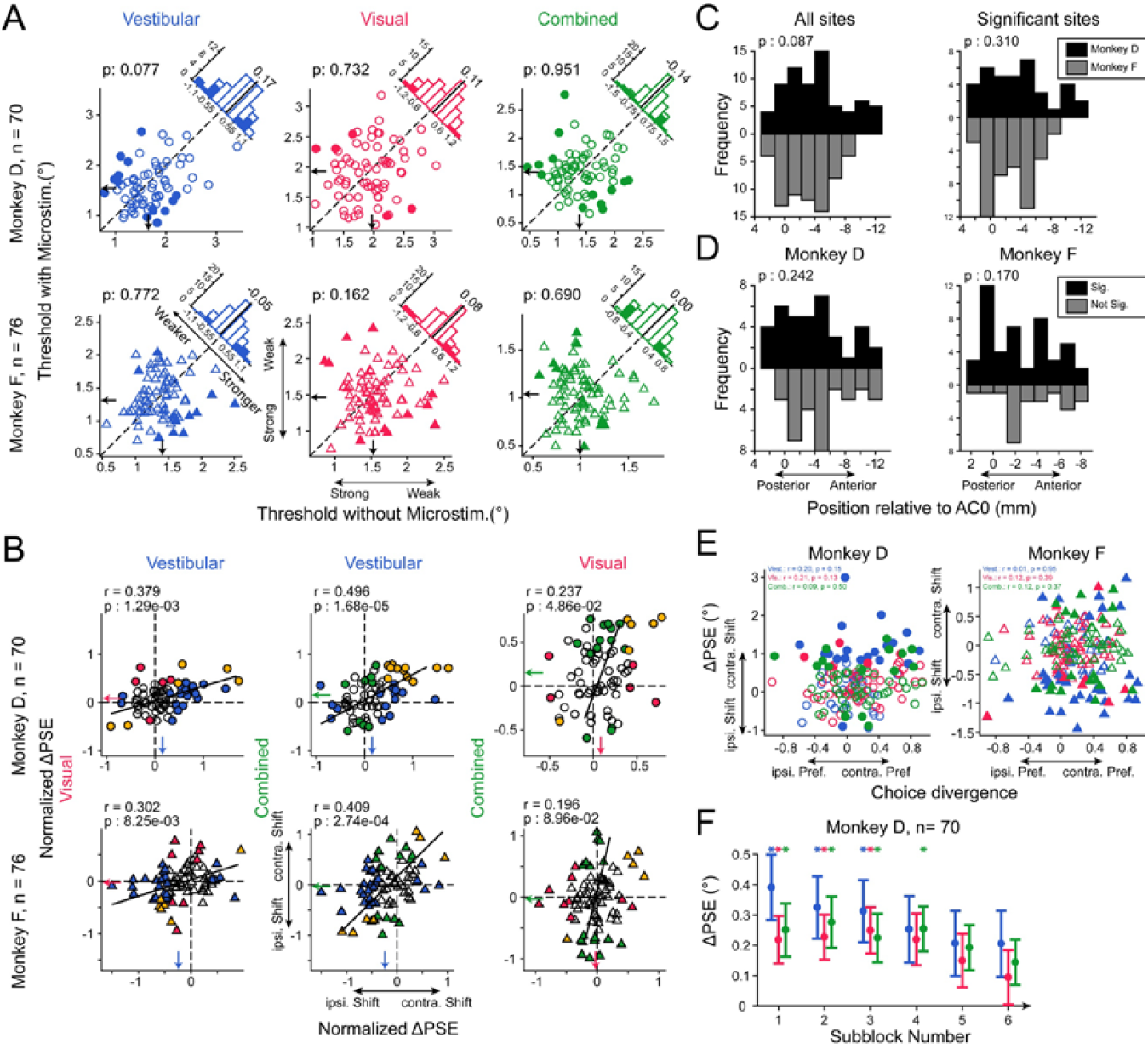
Effects of unilateral microstimulation in CN on perceptual sensitivity and other aspects. **a**, Effects the microstimulation during the stimulus period induces on monkeys’ psychometric thresholds. Labels are same as Fig. 6d. **b**, Cross-modality correlation of microstimulation effects on PSE shift. PSE shift was normalized with respect to perceptual sensitivity (threshold). Colors of dots mean in which cue condition microstimulation significantly biases animal choices (blue, just in vestibular-only condition; red, just in visual-only condition; green, just in combined condition; orange, in both conditions; empty, in neither condition). In each plot, black line was fitted using least sum of squared perpendicular distances; p values and correlation coefficient rare from Spearman correlation analysis. **c-d**, Spatial locations of microstimulation sites across monkeys (**c**) and within individual animal (**d**). In each panel, ordinate and abscissa respectively indicate number of cases and the location relative to AC0 on the AP axis (positive, posterior to AC0; negative, anterior to AC0); color means different monkeys (black, monkey D; gray, monkey F); the p value is from Mann-Whitney U-test. **e**, Correlation between microstimulation effects on PSE (ordinate) and choice preferences (abscissa) of the local multi-units. Filled dots indicate cases with significant microstimulation induced PSE shift; p value and correlation coefficient rare from Spearman correlation. **f**, Adaptation of microstimulation effects. The ΔPSE was calculated in a sliding 5-repetition windows (step is 1 repetition) for all cases from monkey D. Dots with error bars represent mean with SEM. Asterisks mark the subblocks where microstimulation significantly biases PSE (two-tailed t-test; significance level is 0.01).

## Notes

### Competing Interest Statement

The authors have declared no competing interest.

### Summary of Updates

format changed; abstract updated; details added.

## REFERENCES

1. Lee, D., Seo, H. & Jung, M.W. Neural basis of reinforcement learning and decision making. Annu Rev Neurosci 35, 287–308 (2012).

2. Christophel, T.B., Klink, P.C., Spitzer, B., Roelfsema, P.R. & Haynes, J.D. The Distributed Nature of Working Memory. Trends Cogn Sci 21, 111–124 (2017).

3. Steinmetz, N.A., Zatka-Haas, P., Carandini, M. & Harris, K.D. Distributed coding of choice, action and engagement across the mouse brain. Nature 576, 266–273 (2019).

4. Gold, J.I. & Shadlen, M.N. The neural basis of decision making. Annu Rev Neurosci 30, 535–574 (2007).

5. Siegel, M., Buschman, T.J. & Miller, E.K. Cortical information flow during flexible sensorimotor decisions. Science 348, 1352–1355 (2015).

6. Khilkevich, A., et al. Brain-wide dynamics linking sensation to action during decision-making. Nature (2024).

7. Shadlen, M.N. & Newsome, W.T. Neural basis of a perceptual decision in the parietal cortex (area LIP) of the rhesus monkey. J Neurophysiol 86, 1916–1936 (2001).

8. Kim, J.N. & Shadlen, M.N. Neural correlates of a decision in the dorsolateral prefrontal cortex of the macaque. Nat Neurosci 2, 176–185 (1999).

9. Horwitz, G.D., Batista, A.P. & Newsome, W.T. Representation of an abstract perceptual decision in macaque superior colliculus. J Neurophysiol 91, 2281–2296 (2004).

10. Ding, L. & Gold, J.I. Caudate encodes multiple computations for perceptual decisions. J Neurosci 30, 15747–15759 (2010).

11. Li, N., Daie, K., Svoboda, K. & Druckmann, S. Robust neuronal dynamics in premotor cortex during motor planning. Nature 532, 459–464 (2016).

12. Hikosaka, O., Sakamoto, M. & Usui, S. Functional properties of monkey caudate neurons. I. Activities related to saccadic eye movements. J Neurophysiol 61, 780–798 (1989).

13. Kravitz, A.V., et al. Regulation of parkinsonian motor behaviours by optogenetic control of basal ganglia circuitry. Nature 466, 622–626 (2010).

14. Cai, X., Kim, S. & Lee, D. Heterogeneous coding of temporally discounted values in the dorsal and ventral striatum during intertemporal choice. Neuron 69, 170–182 (2011).

15. Kim, Hyoung F. & Hikosaka, O. Distinct Basal Ganglia Circuits Controlling Behaviors Guided by Flexible and Stable Values. Neuron 79, 1001–1010 (2013).

16. Mello, G.B., Soares, S. & Paton, J.J. A scalable population code for time in the striatum. Curr Biol 25, 1113–1122 (2015).

17. Monteiro, T., et al. Using temperature to analyze the neural basis of a time-based decision. Nat Neurosci 26, 1407–1416 (2023).

18. Ding, L. & Gold, J.I. The basal ganglia’s contributions to perceptual decision making. Neuron 79, 640–649 (2013).

19. Lo, C.C. & Wang, X.J. Cortico-basal ganglia circuit mechanism for a decision threshold in reaction time tasks. Nat Neurosci 9, 956–963 (2006).

20. Peters, A.J., Fabre, J.M.J., Steinmetz, N.A., Harris, K.D. & Carandini, M. Striatal activity topographically reflects cortical activity. Nature 591, 420–425 (2021).

21. Chen, S., et al. Brain-wide neural activity underlying memory-guided movement. Cell 187, 676–691 e616 (2024).

22. Van Hoesen, G.W., Yeterian, E.H. & Lavizzo-Mourey, R. Widespread corticostriate projections from temporal cortex of the rhesus monkey. J Comp Neurol 199, 205–219 (1981).

23. Ding, L. & Gold, J.I. Separate, causal roles of the caudate in saccadic choice and execution in a perceptual decision task. Neuron 75, 865–874 (2012).

24. Hanks, T.D., et al. Distinct relationships of parietal and prefrontal cortices to evidence accumulation. Nature 520, 220–223 (2015).

25. Yartsev, M.M., Hanks, T.D., Yoon, A.M. & Brody, C.D. Causal contribution and dynamical encoding in the striatum during evidence accumulation. Elife 7 (2018).

26. Gu, Y., Angelaki, D.E. & Deangelis, G.C. Neural correlates of multisensory cue integration in macaque MSTd. Nat Neurosci 11, 1201–1210 (2008).

27. Hou, H., Zheng, Q., Zhao, Y., Pouget, A. & Gu, Y. Neural Correlates of Optimal Multisensory Decision Making under Time-Varying Reliabilities with an Invariant Linear Probabilistic Population Code. Neuron 104, 1010–1021 e1010 (2019).

28. Zheng, Q., Zhou, L. & Gu, Y. Temporal synchrony effects of optic flow and vestibular inputs on multisensory heading perception. Cell Rep 37, 109999 (2021).

29. Butler, J.S., Smith, S.T., Campos, J.L. & Bulthoff, H.H. Bayesian integration of visual and vestibular signals for heading. J Vis 10, 23 (2010).

30. Friedman, A., et al. A Corticostriatal Path Targeting Striosomes Controls Decision-Making under Conflict. Cell 161, 1320–1333 (2015).

31. Raposo, D., Kaufman, M.T. & Churchland, A.K. A category-free neural population supports evolving demands during decision-making. Nat Neurosci 17, 1784–1792 (2014).

32. Kobak, D., et al. Demixed principal component analysis of neural population data. Elife 5 (2016).

33. Rigotti, M., et al. The importance of mixed selectivity in complex cognitive tasks. Nature 497, 585–590 (2013).

34. Churchland, M.M., et al. Neural population dynamics during reaching. Nature 487, 51–56 (2012).

35. Mante, V., Sussillo, D., Shenoy, K.V. & Newsome, W.T. Context-dependent computation by recurrent dynamics in prefrontal cortex. Nature 503, 78–84 (2013).

36. Drugowitsch, J., DeAngelis, G.C., Klier, E.M., Angelaki, D.E. & Pouget, A. Optimal multisensory decision-making in a reaction-time task. Elife 3 (2014).

37. Bjorklund, A. & Dunnett, S.B. Dopamine neuron systems in the brain: an update. Trends Neurosci 30, 194–202 (2007).

38. Kitama, T., Ohno, T., Tanaka, M., Tsubokawa, H. & Yoshida, K. Stimulation of the caudate nucleus induces contraversive saccadic eye movements as well as head turning in the cat. Neurosci Res 12, 287–292 (1991).

39. Nichols, M.J. & Newsome, W.T. Middle temporal visual area microstimulation influences veridical judgments of motion direction. J Neurosci 22, 9530–9540 (2002).

40. Yu, X., Hou, H., Spillmann, L. & Gu, Y. Causal Evidence of Motion Signals in Macaque Middle Temporal Area Weighted-Pooled for Global Heading Perception. Cereb Cortex 28, 612–624 (2018).

41. Histed, M.H., Bonin, V. & Reid, R.C. Direct activation of sparse, distributed populations of cortical neurons by electrical microstimulation. Neuron 63, 508–522 (2009).

42. Tolias, A.S., et al. Mapping cortical activity elicited with electrical microstimulation using FMRI in the macaque. Neuron 48, 901–911 (2005).

43. Jeurissen, D., Shushruth, S., El-Shamayleh, Y., Horwitz, G.D. & Shadlen, M.N. Deficits in decision-making induced by parietal cortex inactivation are compensated at two timescales. Neuron 110, 1924–1931 e1925 (2022).

44. Gu, Y., Deangelis, G.C. & Angelaki, D.E. Causal links between dorsal medial superior temporal area neurons and multisensory heading perception. J Neurosci 32, 2299–2313 (2012).

45. Potegal, M., Copack, P., de Jong, J., Krauthamer, G. & Gilman, S. Vestibular input to the caudate nucleus. Exp Neurol 32, 448–465 (1971).

46. Maunsell, J.H.R. & Vanessen, D.C. The Connections of the Middle Temporal Visual Area (Mt) and Their Relationship to a Cortical Hierarchy in the Macaque Monkey. J Neurosci 3, 2563–2586 (1983).

47. Reig, R. & Silberberg, G. Multisensory integration in the mouse striatum. Neuron 83, 1200–1212 (2014).

48. Borra, E., Biancheri, D., Rizzo, M., Leonardi, F. & Luppino, G. Crossed Corticostriatal Projections in the Macaque Brain. J Neurosci 42, 7060–7076 (2022).

49. Fetsch, C.R., Turner, A.H., DeAngelis, G.C. & Angelaki, D.E. Dynamic reweighting of visual and vestibular cues during self-motion perception. J Neurosci 29, 15601–15612 (2009).

50. Nienborg, H. & Cumming, B.G. Decision-related activity in sensory neurons reflects more than a neuron’s causal effect. Nature 459, 89–92 (2009).

51. Yates, J.L., Park, I.M., Katz, L.N., Pillow, J.W. & Huk, A.C. Functional dissection of signal and noise in MT and LIP during decision-making. Nat Neurosci 20, 1285–1292 (2017).

52. Okazawa, G., Sha, L., Purcell, B.A. & Kiani, R. Psychophysical reverse correlation reflects both sensory and decision-making processes. Nat Commun 9, 3479 (2018).

53. Noudoost, B. & Moore, T. Control of visual cortical signals by prefrontal dopamine. Nature 474, 372–375 (2011).

54. Ott, T., Jacob, S.N. & Nieder, A. Dopamine receptors differentially enhance rule coding in primate prefrontal cortex neurons. Neuron 84, 1317–1328 (2014).

55. Puig, M.V. & Miller, E.K. The role of prefrontal dopamine D1 receptors in the neural mechanisms of associative learning. Neuron 74, 874–886 (2012).

56. Williams, Z.M. & Eskandar, E.N. Selective enhancement of associative learning by microstimulation of the anterior caudate. Nat Neurosci 9, 562–568 (2006).

57. Nicola, S.M., Surmeier, J. & Malenka, R.C. Dopaminergic modulation of neuronal excitability in the striatum and nucleus accumbens. Annu Rev Neurosci 23, 185–215 (2000).

58. Vijayraghavan, S., Wang, M., Birnbaum, S.G., Williams, G.V. & Arnsten, A.F. Inverted-U dopamine D1 receptor actions on prefrontal neurons engaged in working memory. Nat Neurosci 10, 376–384 (2007).

59. Bolkan, S.S., et al. Opponent control of behavior by dorsomedial striatal pathways depends on task demands and internal state. Nat Neurosci 25, 345–357 (2022).

60. Chen, Y., et al. Circuit-specific gene therapy reverses core symptoms in a primate Parkinson’s disease model. Cell 186, 5394–5410 e5318 (2023).

61. Hanks, T.D., Ditterich, J. & Shadlen, M.N. Microstimulation of macaque area LIP affects decision-making in a motion discrimination task. Nat Neurosci 9, 682–689 (2006).

62. Salzman, C.D., Murasugi, C.M., Britten, K.H. & Newsome, W.T. Microstimulation in visual area MT: effects on direction discrimination performance. J Neurosci 12, 2331–2355 (1992).

63. Doi, T., Fan, Y., Gold, J.I. & Ding, L. The caudate nucleus contributes causally to decisions that balance reward and uncertain visual information. Elife 9 (2020).

64. Roitman, J.D. & Shadlen, M.N. Response of neurons in the lateral intraparietal area during a combined visual discrimination reaction time task. J Neurosci 22, 9475–9489 (2002).

65. Gu, Y., Watkins, P.V., Angelaki, D.E. & DeAngelis, G.C. Visual and nonvisual contributions to three-dimensional heading selectivity in the medial superior temporal area. J Neurosci 26, 73–85 (2006).

66. Britten, K.H., Shadlen, M.N., Newsome, W.T. & Movshon, J.A. The analysis of visual motion: a comparison of neuronal and psychophysical performance. J Neurosci 12, 4745–4765 (1992).

67. Hikosaka, O., Sakamoto, M. & Usui, S. Functional properties of monkey caudate neurons. III. Activities related to expectation of target and reward. J Neurophysiol 61, 814–832 (1989).

68. Lanciego, J.L., Luquin, N. & Obeso, J.A. Functional neuroanatomy of the basal ganglia. Cold Spring Harb Perspect Med 2, a009621 (2012).

69. Li, W., Lu, J., Zhu, Z. & Gu, Y. Causal contribution of optic flow signal in Macaque extrastriate visual cortex for roll perception. Nat Commun 13, 5479 (2022).

70. Watanabe, M. & Munoz, D.P. Saccade reaction times are influenced by caudate microstimulation following and prior to visual stimulus appearance. J Cogn Neurosci 23, 1794–1807 (2011).

